# Defining epidermal basal cell states during skin homeostasis and wound healing using single-cell transcriptomics

**DOI:** 10.1101/793117

**Authors:** Daniel Haensel, Suoqin Jin, Rachel Cinco, Peng Sun, Quy Nguyen, Zixuan Cang, Morgan Dragan, Yanwen Gong, Adam L MacLean, Kai Kessenbrock, Enrico Gratton, Qing Nie, Xing Dai

## Abstract

Our knowledge of transcriptional heterogeneities in epithelial stem/progenitor cell compartments is limited. Epidermal basal cells sustain cutaneous tissue maintenance and drive wound healing. Previous studies have probed basal cell heterogeneity in stem/progenitor potential, but a non-biased dissection of basal cell dynamics during differentiation is lacking. Using single-cell RNA-sequencing coupled with RNAScope and fluorescence lifetime imaging, we identify three non-proliferative and one proliferative basal cell transcriptional states in homeostatic skin that differ in metabolic preference and become spatially partitioned during wound re-epithelialization. Pseudotemporal trajectory and RNA velocity analyses produce a quasi-linear differentiation hierarchy where basal cells progress from *Col17a*1^high^/*Trp63*^high^ state to early response state, proliferate at the juncture of these two states, or become growth arrested before differentiating into spinous cells. Wound healing induces plasticity manifested by dynamic basal-spinous interconversions at multiple basal states. Our study provides a systematic view of epidermal cellular dynamics supporting a revised “hierarchical-lineage” model of homeostasis.

## INTRODUCTION

Epithelial tissue maintenance is driven by resident stem cells, the proliferation and differentiation dynamics of which need to be tailored to the tissue’s homeostatic and regenerative needs. However, our understanding of tissue-specific cellular dynamics *in vivo* at both single-cell and tissue scales is often very limited. The self-renewing skin epidermis represents an outstanding model to study the precise sequence of events that underlie the commitment and differentiation of epithelial stem cells towards highly specialized terminal states with important biological functions.

Within the adult mouse interfollicular epidermis, stem/progenitor cells residing in the basal layer undergo self-renewing or differentiative cell divisions to maintain a proper pool of basal cells and to generate post-mitotic differentiating (spinous, granular) cells in the suprabasal layers that ultimately form the stratum corneum – an outer permeability barrier that protects an organism from dehydration, infection, and a myriad of other harmful insults (Gonzales and Fuchs, 2017). Cumulative evidence supports multiple possible mechanisms of epidermal homeostasis: 1) a hierarchical lineage of relatively quiescent stem cells giving rise to faster cycling, committed progenitor cells that then exit the cell cycle and terminally differentiate; 2) a single, equipotent population of progenitor cells stochastically choosing between self-renewal and differentiation; and 3) two spatially segregated populations of stem cells that divide at different rates and adopt distinct lineage trajectories (Gonzales and Fuchs, 2017; Mascré et al., 2012; Rompolas et al., 2016; Sada et al., 2016). The different criteria used for stem/progenitor fate assignment (often linked to the specific techniques used), such as molecular differentiation markers, basal layer residence status, and assumptions about stem-cell-division or clonal-growth kinetics, may account for the differences in data interpretation (Gonzales and Fuchs, 2017). Moreover, the observed epidermal stem cell heterogeneity in mouse back skin may reflect different cellular states of a single differentiation program (Rognoni and Watt, 2018). Clearly, genome-wide transcriptomic and *in situ* analyses with single-cell resolution are needed to provide a comprehensive, non-biased picture of basal cell heterogeneity, to unravel previously unknown cellular states and their transitions during epidermal lineage differentiation, and to seek unifying principles of epidermal homeostasis.

Upon cutaneous wounding, the skin must alter its cellular dynamics to facilitate efficient healing for timely restoration of the protective barrier. Wound healing represents a highly regulated process composed of several distinct but overlapping stages (inflammation, re-epithelialization, and resolution) that involve the coordinated activities of epidermal, dermal, immune and endothelial cells (Gurtner et al., 2008). Re-epithelialization is driven by spatially patterned migration and proliferation of epidermal cells at the wound periphery, as well as migration and dedifferentiation/reprogramming of hair follicle (HF) and sebaceous gland epithelial cells (Haensel and Dai, 2018; Park et al., 2017; Rognoni and Watt, 2018). What and how epidermal cells migrate during wound re-epithelialization were a subject of debate, with two different models being proposed in the past: 1) basal cells first migrate into the wound bed and unidirectionally convert into suprabasal cells; and 2) wound peripheral epidermal cells crawl or “leapfrog” over one another such that suprabasal cells migrate in and become basal cells (Rittié, 2016; Rognoni and Watt, 2018). Recent live cell imaging and lineage tracing studies have provided clarity to this issue by defining distinct zones of epidermal cellular activities in the wound area: a migratory zone next to the wound margin where both basal and suprabasal cells move towards the wound center, an intermediate, mixed zone of coordinated migration and proliferation, and a hyperproliferative zone furthest away from the wound margin (Aragona et al., 2017; Park et al., 2017). Hrecisely how many distinct transcriptional states exist for wound epidermal cells and whether these states correlate with or differ from their homeostatic counterparts, particularly within the basal layer, remains to be elucidated.

In this work, we performed single-cell RNA-sequencing (scRNA-Seq) analysis on thousands of cells derived from normal mouse back skin or wounded skin at the re-epithelization stage. Our analysis identifies four distinct states of basal cells in normal skin that shift proportions and gene expression during wound healing. Using multiplexed RNA *in situ* detection (RNAScope) and fluorescence lifetime imaging microscopy (FLIM), we spatially map the scRNA-Seq-revealed molecular and metabolic heterogeneities onto the intact normal and wounded skin tissue. Using standard and custom tools to predict cell trajectories and lineage relationships, we place the different basal cell states temporally onto a differentiation hierarchy and identify enhanced cell fate fluidity featuring increased basal-spinous interconversions during wound healing. Overall, our study provides a comprehensive single-cell perspective of epidermal cellular dynamics and transitional states during normal homeostasis and repair that is consistent with the “hierarchical-lineage” model of epidermal homeostasis but encompassing more than one possible stem/progenitor cell states.

## RESULTS

### scRNA-Seq reveals global changes in skin cellular makeup during wound healing

To systematically examine major cell type and cell state differences between skin homeostasis and repair, we performed scRNA-Seq on samples isolated from unwounded (UW) and wounded (WO) mouse back skin (Figure 1A). The wound samples were taken at 4 days after the introduction of 6-mm wounds, corresponding to a stage of active re-epithelialization (Figure 1B and S1A). We performed read depth normalization, visualized control metrics (Figure S1B; see Methods section), and obtained a total of 10,615 (from 2 UW biological replicates) and 16,164 (from 3 WO biological replicates) cells for downstream analyses. When data from all 5 samples were combined, cells belonging to each major cell type – epithelial (identified by *Krt14* and *Krt1* expression), fibroblast (identified by *Col1a2* expression), and immune (identified by *Cd45* expression) - segregated together in the overall tSNE plot, and the immune cells showed greater changes between UW and WO samples than fibroblasts and epithelial cells (Figure 1C, 1D, and S2). Moreover, the relative percentages of immune cells and fibroblasts were increased in the WO samples at the expense of epithelial cells (Figure 1E).

**Figure 1:**
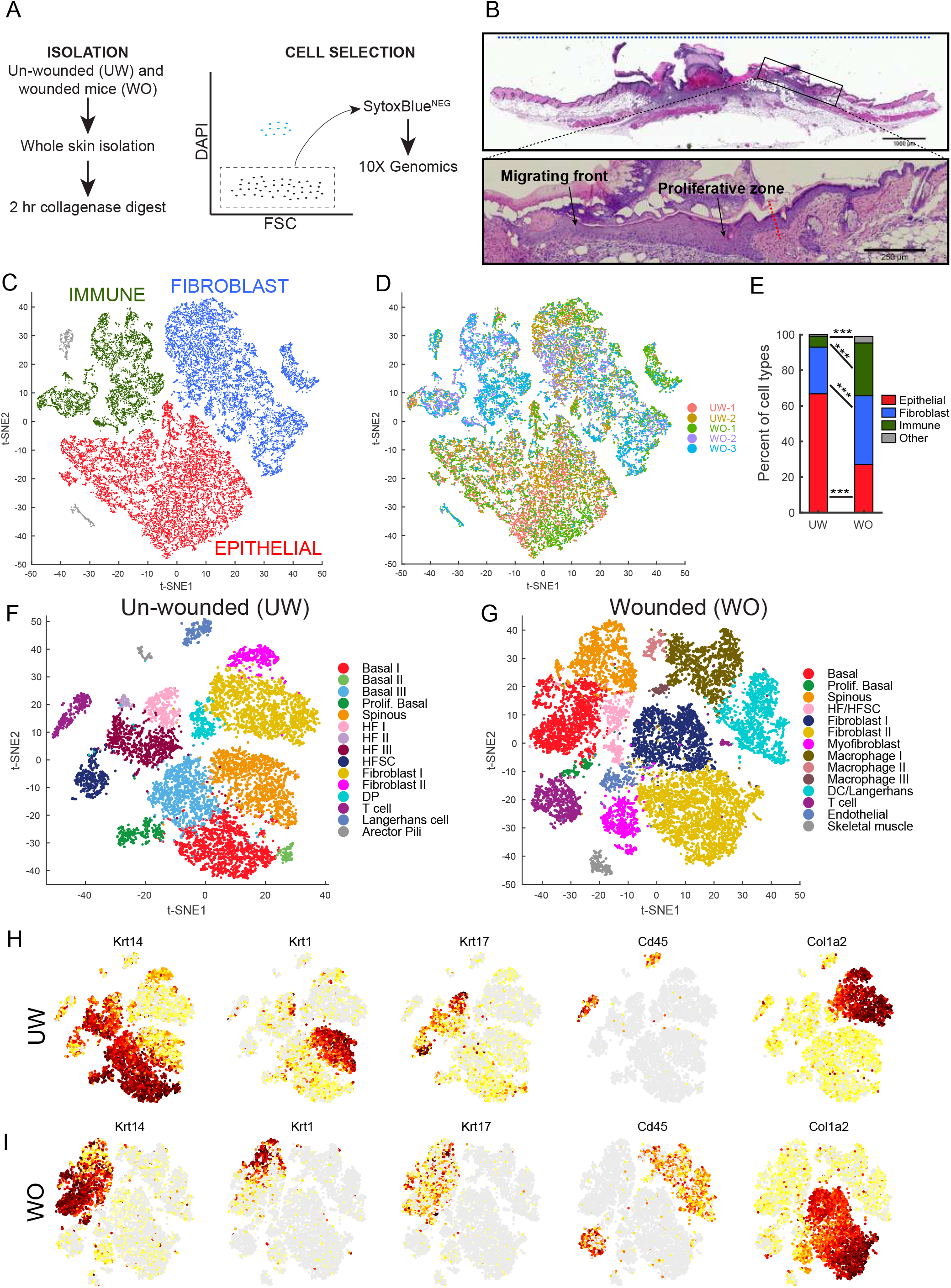
scRNA-Seq data on all cell types of the UW and WO skin. A. Schematic diagram detailing the single cell isolation and live cell selection strategy. B. H/E analysis of WO skin showing a region equivalent to those used for scRNA-Seq. Blue dashed line indicates representative 10 mm region used for single cell suspension. Enlarged image of the boxed area in top panel is shown at the bottom to highlight the wound migrating front and proliferative zone. Red dashed line indicates wound margin. C. tSNE plot for all samples (2 UW and 3 WO) with the major cell type populations (epithelial, fibroblast, and immune) highlighted. D. tSNE plot for all samples (2 UW and 3 WO) with each replicate sample ID identified by a unique color identifier. E. Bar graph representing major cell type populations in the UW and WO samples. Chi-squared test was used to determine whether there were significant proportion differences of each cell type between UW and WO samples. *** *p* < 0.0005. F. tSNE plot for the 2 UW replicate datasets that were aggregated and then batch corrected using CCA. G. tSNE plot for the 3 WO replicate datasets that were aggregated and then batch corrected using CCA. Principle components and resolution parameters utilized in (C) were also utilized in (D). H. Feature plots of the indicated genes in UW sample. I. Feature plots of the indicated genes in WO sample.

We also analyzed the UW and WO sample types separately, surveying major cellular compositions of the skin under each condition. Biological replicates were aggregated together, and batch corrected using canonical correlation analysis (CCA) implemented in the Seurat R package (Butler et al., 2018) (Figure S3A and S3B). Marker genes were found for each individual t-distributed stochastic neighbor embedding (tSNE) cluster and used in tandem with known cell type markers (Han et al., 2018; Joost et al., 2016, 2018; Meador et al., 2014) to determine the identity of each cluster (Figure 1F-G, S3C, and S3D; Table S1 and S2). Using identical parameters for UW and WO samples, we observed 15 and 14 cell clusters, respectively (Figure 1F and 1G). Feature plots of key cell type markers revealed population-level changes in epidermal basal cells (*Krt14*), epidermal spinous cells (*Krt1*), HF-associated cells (*Krt17*), immune cells (*Cd45*), and fibroblasts (*Col1a2*) (Figure 1H and 1I). Several new cell types, including macrophages, dendritic cells (DC, which also includes the Cd207^+^ Langerhans cells), myofibroblasts, and endothelial cells, were detected only in the WO skin.

Overall, these data provide the first general overview of the major changes in cellular heterogeneity from homeostasis to a wound healing state, and they support existing knowledge obtained using traditional methods (Gurtner et al., 2008; Shaw and Martin, 2009).

### Wound epidermal basal cells upregulate inflammation- and migration-related genes expression

Epithelial cells of the interfollicular epidermis and its associated appendages (HF, sebaceous gland) drive wound re-epithelialization. When these cells were subset out from the 5 sample-aggregate data, comparable numbers of epidermal and HF cell clusters were observed in the UW and WO samples (Figure S4 and S5; Table S3 and S4). Both UW and WO samples contained 4 basal cell subclusters, 2 spinous cell subclusters, and 4 HF subclusters, which are generally consistent with the reported scRNA-Seq data using Fludigm C1 platform to sequence 1422 cells (Joost et al., 2016). Compared to the *Joost et al.* dataset, our dataset does not contain a minor loricrin^+^ granular cell population but features a distinct proliferative basal cell subcluster in both UW and WO skin (Figure 1F, 1G, S4A, S4B, S4E, and S4F). Overall, the relative proportions of the various epithelial cell types did not dramatically change between UW and WO skin (Figure S4G).

Published scRNA-Seq data on wounded skin focused on HF-resident Lgr5- or Lgr6-expressing stem cells and their progenies (Joost et al., 2018). As such, little is known about the single-cell transcriptomic dynamics of interfollicular epidermal basal cells that are major players in wound re-epithelialization. To this end, we first compared UW and WO basal cells by subseting out the *Krt14*^+^, but *Krt17^-^CD34^-^ (*HF markers), cells from the epithelial subsets (Figure S4E and S4F). We also excluded the proliferative *Krt14*^+^ basal cell subcluster from this analysis to avoid cell cycle gene expression overshadowing other molecular differences (Figure S6A-C). A total of 53 and 99 genes were particularly enriched in UW and WO basal cells, respectively (Table S5). Interestingly, the expression levels of such enriched genes were not uniform across all single cells of each condition, with some UW basal cells displaying a WO-like signature but not vice versa (Figure 2A).

**Figure 2:**
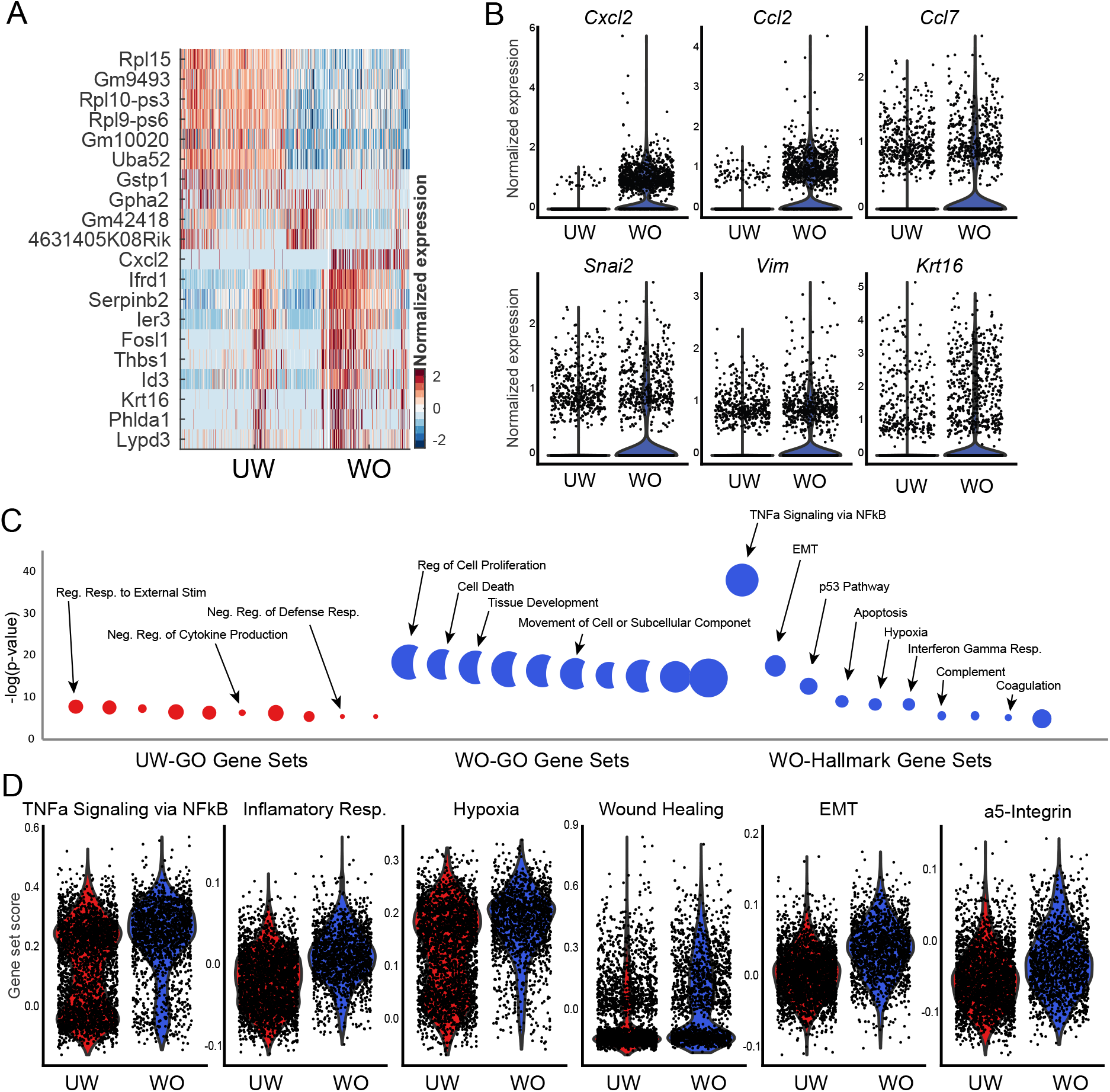
scRNA-Seq data comparing gene expression in bulk epidermal basal cells of the UW and WO skin. A. Heatmap showing the top 10 markers for basal cells from the UW and WO samples. B. Violin plots showing expression of select genes for UW and WO basal cells. C. GO analysis of basal cells from UW and WO samples using two different gene sets. Hallmark Gene Sets are defined by fewer genes than GO Gene Sets. D. Gene scoring analysis of UW and WO basal cells using the indicated molecular signatures.

Genes upregulated in WO basal cells included inflammatory genes *Cxcl2*, *Ccl2,* and *Ccl7*, epithelial-to-mesenchymal transition (EMT)-related genes *Snai2* and *Vim*, and positive control *Krt16* (Wawersik et al., 2013) (Figure 2B). Gene Ontology (GO) analysis of all differentially expressed genes revealed enrichment of inflammation (e.g., TNF-α signaling, interferon gamma response) and EMT signatures in WO basal cells (Figure 2C). Comparing UW and WO basal cells for specific gene sets showed upregulation of genes associated with wound healing, EMT, and those that are up-regulated in the migratory front of wound neo-epidermis (i.e., α5 integrin-expressing cells) (Aragona et al., 2017) (Figure 2D). Interestingly, UW basal cells encompassed two distinct subsets scoring low and high for TNF-α signaling, inflammatory response, and hypoxia, whereas the WO basal cells scored uniformly higher for the same gene sets (Figure 2D). Together, these data demonstrate that epidermal basal cells exist in distinct inflammation-low and inflammation-high states during homeostasis, and that they dramatically upregulate inflammatory and migratory gene expression during wound healing.

### Three distinct non-proliferative basal cell states exist in un-wounded skin

We next zoomed in on the non-proliferative UW basal cells in order to delineate their unknown heterogeneity in an unbiased fashion. Indeed, three distinct tSNE subclusters were observed (Figure 3A), and each was enriched for specific marker genes (Figure 3B; Table S6). Subcluster 1 was defined by top markers such as *Col17a1* and *Trp63*, and was therefore named the *Col17a1*^Hi^ subcluster (Figure 3C, S7A, and S7B; Table S6). *Col17a1* encodes a hemidesmosome component that negatively regulates epidermal proliferation (Watanabe et al., 2017), and its high expression marks a subset of symmetrically-dividing epidermal cells with growth advantage over the asymmetrically-dividing *Col17a1*^low^ cells (Liu et al., 2019). *Trp63* encodes a transcription factor (TF) with master-regulatory role in epidermal development and adult epidermal homeostasis (Sano et al., 2007; Truong et al., 2006), and is expressed at a higher level in quiescent bulge HF stem cells (Bu-HFSCs) than the activated stem/progenitor cells (Lien et al., 2011). Subcluster 2 was named the early response (ER) subcluster due to enrichment for a number of immediate early genes (Figure 3C and S7A; Table S6) (Herschman, 1991). Top markers of this subcluster also include those associated with activated Bu-HFSCs relative to quiescent Bu-HFSCs (Lien et al., 2011) or with known function in regulating proliferation (Andrianne et al., 2017; Briso et al., 2013; Florin et al., 2006; Rotzer et al., 2006; Zenz and Wagner, 2006; Zhu et al., 2008) (Figure 3C and S7A; Table S6). Subcluster 3 was designated the growth arrested (GA) subcluster because of enriched expression of genes with known functions in promoting cell cycle arrest, such as *Cdkn1a*, *Irf6*, *Ovol1*, and *Sfn* (Hammond et al., 2012; Ingraham et al., 2006; Nair et al., 2006; Topley et al., 1999) (Figure 3C and S7A; Table S6). GA- and *Col17a1*^Hi^-enriched genes presented a trend of inverse correlation among the 3 subclusters (Figure 3C). Together, these data show the existence of 3 major, distinct transcriptional states in non-proliferative UW basal cells.

**Figure 3:**
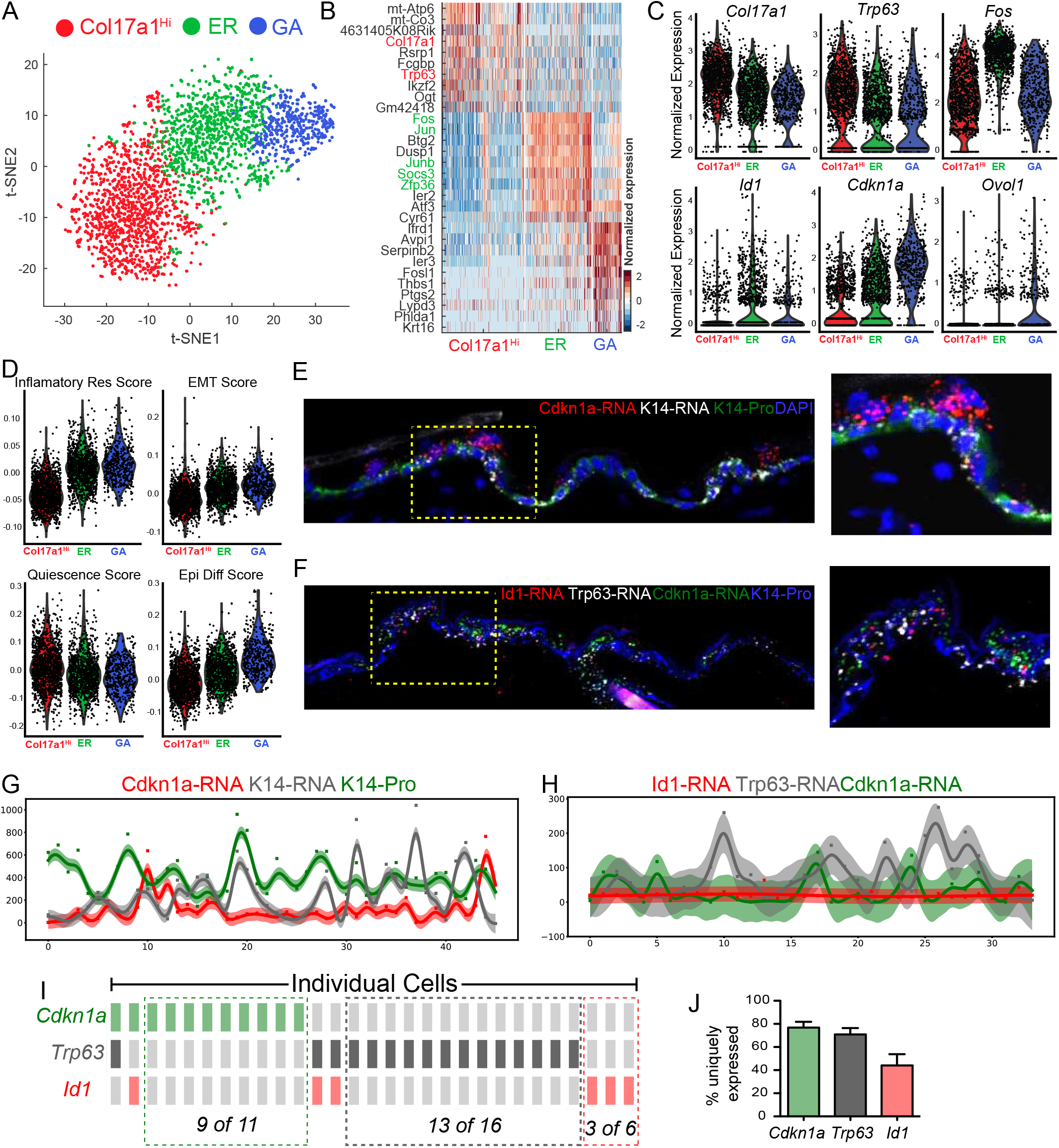
scRNA-Seq and RNAScope data revealing heterogeneity within UW epidermal basal cells. A. tSNE plot for non-proliferative basal cells from UW sample. B. Heatmap of top 10 markers for the different basal cell subclusters in (A). Genes used to annotate cell identity are indicated by the corresponding colors. C. Violin plots of select key marker genes in the non-proliferative basal subclusters from UW sample. D. Gene scoring analysis of the different non-proliferative basal subclusters from UW sample using the indicated molecular signatures. E. RNAScope data showing spatial distribution of *Cdkn1a*, *Krt14* transcripts and K14 protein in UW skin. DAPI stains the nuclei. Enlarged image of the boxed area in left panel is shown on the right. F. RNAScope data showing spatial distribution of *Cdkn1a*, *Trp63*, *Id1* transcripts and K14 protein in UW skin. Enlarged image of the boxed area in left panel is shown on the right. G. Quantification of fluorescence intensity (represented by a color-coded dot) for *Cdkn1a*, *Krt14* transcripts and K14 protein in each individual cell from a representative section. The curve represents a Gaussian Process Regression (GPR) and a 95% confidence interval is shown as shaded area. H. Quantification of fluorescence intensity for *Cdkn1a*, *Trp63*, *Id1* transcripts and K14 protein in each individual cell from a representative section. See legends for (G). I. OncoPrint representation of *Cdkn1a*, *Trp63*, and *Id1* expression in individual cells where a column represents an individual cell. A color-coded rectangle indicates high expression of the corresponding marker gene. The cells are sorted based on the on/off state of the markers to show mutually exclusive expression pattern. J. Bar graph showing percent cells that exclusively express *Cdkn1a*, *Trp63*, or *Id1* of the total number of cells expressing the particular gene (n = 3 replicates). Error bars represent mean +/− SEM.

The 3 non-proliferative basal cell states also showed differential expression of inflammation- and EMT-related genes. The GA state showed the highest expression of TNFα signaling, hypoxia, EMT genes, and it was also enriched for a gene signature derived from a label-retaining basal cell (LRC) population (Sada et al., 2016) (Figure 3D, S7C, and S7D). In contrast, the *Col17a1*^Hi^ state scored the lowest for inflammation and EMT genes, but highest for a quiescence signature derived from quiescent stem cells of multiple tissues (Cheung and Rando, 2013) (Figure 3D). There was a step-wise increase in epidermal differentiation gene expression from *Col17a1*^Hi^ to ER and to GA states (Figure 3D). These data suggest that the 3 non-proliferative basal cell states are defined by their differences in inflammation, migration, quiescence, cell cycle exit, and differentiation status.

To validate the existence of 3 non-proliferative basal cell states in the intact skin tissue, we utilized RNAScope, a multiplexed, semi-quantitative *in situ* RNA detection method. Co-analysis of *Cdkn1a* (GA state marker) and *Krt14* transcripts with K14 protein revealed several interesting points: 1) the levels of *Krt14* transcript and to a lesser extent, K14 protein fluctuated along the basal layer of the epidermis, and the location of such variation did not always coincide; 2) *Cdkn1a* transcripts were present in some but not all K14-positive basal cells, and the highest expression was detected in a subset of suprabasal cells (Figure 3E, 3G, S7E). Co-analysis of *Cdkn1a* with *Col17a1*^Hi^ state marker *Trp63* and ER state marker *Id1* revealed cells in the basal layer that uniquely express each of the markers (Figure 3F, 3H-J, S7F, and S7G). Few cells expressed 2 markers simultaneously, *Trp63* and *Cdkn1a* expression is mutually exclusive, and none expressed all 3 markers (Figure 3I and 3J). These results confirm that the epidermal basal cells exist in multiple non-proliferative transcriptional states *in vivo* that are defined by markers identified in our scRNA-Seq analysis.

### Altered proportion and spatial partitioning of the three distinct non-proliferative basal cell states in wounded skin

We next turned to the WO samples to assess basal cell heterogeneity during wound healing. Similar to the UW basal cells, unsupervised analysis indicated the presence of three distinct non-proliferative basal cell subclusters (Figure 4A). By comparing marker genes of the WO vs. UW basal cell subclusters and using Random Forest classification (see Methods section), we found the WO basal cell subclusters to approximately correspond to *Col17a1*^Hi^, ER and GA subclusters in the UW samples (Figure 4B, 4C, and S8A-C; Table S6 and S7). Similar to UW GA cells, the WO GA cells were enriched for genes associated with α5-integrin-positive migrating front, inflammation, hypoxia, and have the lowest quiescence score but the highest epidermal differentiation score (Figure 4E and S9A; Table S7). Moreover, individual genes that are previously known to be enriched (*Krt16*, *Hbegf*, and *Snai2*) and downregulated (*Cd9*) in the wound migratory front (Haensel and Dai, 2018; Jiang et al., 2013; Shirakata, 2005) are enriched and de-enriched, respectively, in the WO GA subcluster (Figure S9B, S9C). These data suggest that cells at the migrating front are predominantly in a GA state (see below). While the UW *Col17a1*^Hi^ state displayed no enrichment for any particular GO term, its WO counterpart showed a significant enrichment for oxidative phosphorylation genes (Figure S9A), a point we will return to later. Overall, compared to the UW skin, the GA subcluster is expanded in the WO skin at the expense of the *Col17a1*^Hi^ subcluster (Figure 4A and 4D).

**Figure 4:**
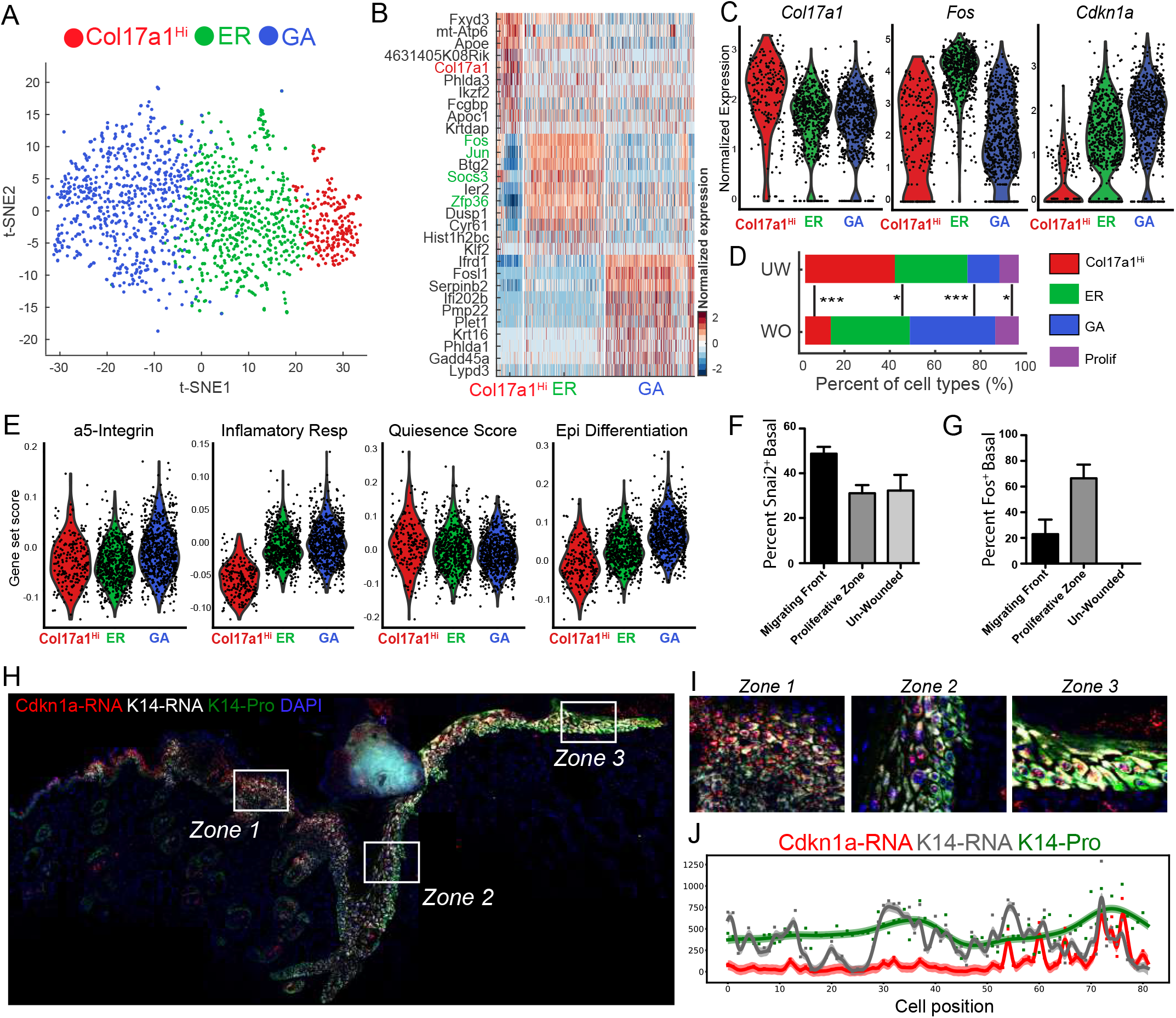
scRNA-Seq and RNAScope data revealing basal cell heterogeneity in WO skin. A. tSNE plot for non-proliferative basal cells from WO sample. B. Heatmap of top 10 markers for the different basal cell subclusters in (a). Genes used to annotate cell identity are indicated by the corresponding colors. C. Violin plots of select key marker genes in the non-proliferative basal subclusters from WO sample. D. Subset composition of UW and WO proliferative and non-proliferative basal cells. Chi-squared test was used to determine statistical significance. \*\*\**p* < 0.0005. \**p* < 0.05. E. Gene scoring analysis of the different basal subclusters from WO sample using the indicated molecular signatures. F. Quantification of percent Snai2^+^ cells in the basal layer of the indicated regions in WO skin. G. Quantification of percent Fos^+^ cells in the basal layer of the indicated regions in WO skin. H. RNAScope data showing spatial distribution of *Cdkn1a*, *Krt14* transcripts and K14 protein in WO skin. DAPI stains the nuclei. I. Enlarged images of the boxed areas in (H). J. Quantification of fluorescence intensity for *Cdkn1a*, *Krt14* transcripts and K14 protein in each individual cell from a representative WO skin section. See Figure 3G legends for additional information.

We next performed immunofluorescence and RNAScope to examine the spatial distribution of basal cell states in WO skin. GA state marker Snai2 (Slug) protein was found to be enriched in basal cells at the migrating front relative to those in the hyperproliferative zone or distal to the wound (Figure 4F and S9D). ER state marker Fos protein was found to be enriched in the proliferative zone, but its expression dissipated in the migratory zone or non-wounded area (Figure 4G, S9E). In the wound hyperproliferative zone, GA state marker *Cdkn1a* was predominantly expressed in suprabasal cells; however, in the migrating front, its strong expression was detected in both basal and suprabasal cells (Figure 4H-J). Quantification of *Cdkn1a* signals along the entire basal layer of the wounded area revealed a clear increase towards the migrating front (Figure 4J).

Collectively, our data show that basal cells in the wounded skin also exist in three major distinct states, with the proliferative zone predominantly defined as the early response state, and the migrating front predominantly defined as the growth arrested state.

### Metabolic heterogeneity in basal cells of the normal and wounded skin

Energy metabolism is fundamentally important in shaping myriad cellular processes during homeostasis and repair, and single-cell analysis enables the dissection of metabolic heterogeneity in normal and wounded skin. To compare the metabolic status of the different basal cell states identified above, we scored all UW and WO basal cells for their expression oxidative phosphorylation and glycolysis genes. In UW skin, the proliferative basal cell states showed the highest, whereas the GA state showed the lowest, oxidative phosphorylation score (Figure 5A). In WO skin, oxidative phosphorylation of the *Col17a1*^Hi^ state was elevated to a level similar to that of proliferative WO basal cells (Figure 5B). The WO GA state still scored the lowest for oxidative phosphorylation, which is in keeping with its highest expression of hypoxia response factor *Hif1a* (Figure 5B and S10B). An apparently opposite trend was seen for glycolysis genes, as GA cells displayed the highest glycolysis score than the other non-proliferative subsets in both UW and WO skin (Figure S10C).

**Figure 5:**
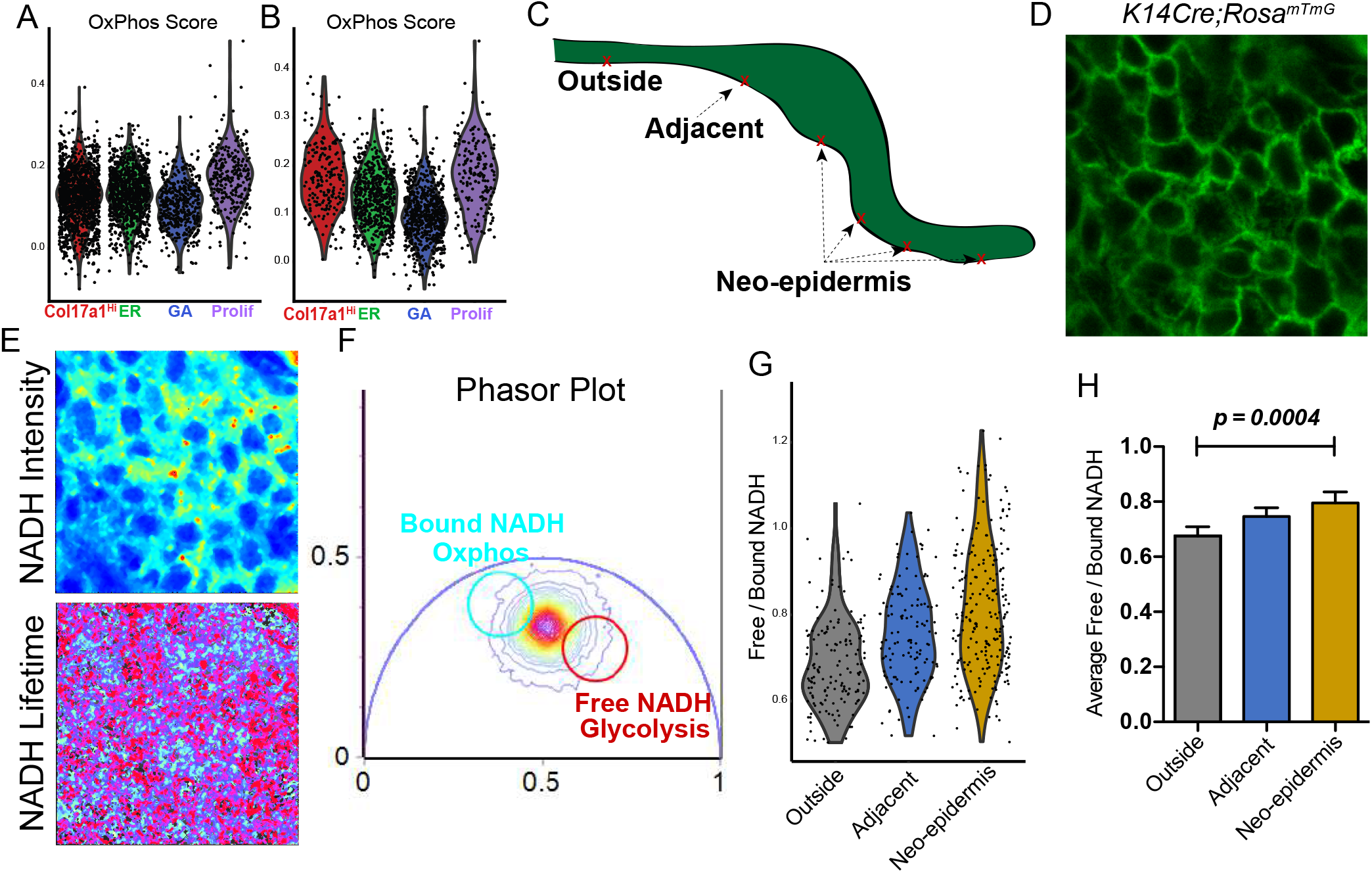
FLIM data validating scRNA-Seq-predicted metabolic heterogeneity in WO skin. A. Gene scoring analysis of all 4 basal subclusters from UW sample using an oxidative phosphorylation signature. B. Gene scoring analysis of all 4 basal subclusters from WO sample using an oxidative phosphorylation signature. C. Sketch diagram of wound epithelium showing the areas probed with FLIM. D. A representative image of wound epidermal cells indicated by GFP expression. E. Representative images of NADH signal and NADH lifetime signal from c. F. A representative phasor plot with cell phasor fingerprint, which is a representation of the fluorescence lifetime decay of all cells in the region of interest (ROI) after fast Fourier transformation. G. Violin plot incorporating all cells and their corresponding free/bound ratios from 4 biological replicates (156 cells from outside region, 127 cells from the adjacent region, and 231 cells from neo-epidermis). H. Quantification of average free/bound NADH ratios from multiple cells from 4 biological replicates of various regions of the wound. For statistical analysis we used an unpaired two-tailed Student’s t-test. Error bars represent mean +/− SEM.

Two-photon excitation (TPE) and NADH fluorescence lifetime imaging (FLIM) have been used for *in vivo* metabolic measurements during skin wound healing (Deka et al., 2016; Jones et al., 2018), but single-cell variation of metabolic activity remains to be assessed. We applied this method on wounded skin derived from *K14-Cre/mTmG* mice, where epidermal cells are visualized by GFP expression *ex vivo*. Care was taken to measure NADH autofluorescence in individual cells (Stringari et al., 2015) within the GFP^+^ basal layer in three areas: outside of the wound, portions of the proliferative zone, and regions of the migratory front deep within the wound (Figure 5C and 5D). TPE NADH intensities and lifetimes were captured and displayed in phasor plots (Figure 5E and 5F). Free-to-bound NADH ratios, indicative of the relative level of oxidative phosphorylation (Stringari et al., 2012, 2015), were calculated from each basal cell in each region. A gourd-shaped distribution of the ratios was observed for cells in the region far from and outside of the wound (Figure 5G), suggesting that basal cells in UW skin can be classified into OxPhos^high^ (less abundant) and OxPhos^low^ states. The ratios were elevated in basal cells immediately adjacent to and within the wound, with the highest values detected in a significant fraction of basal cells in the neo-epidermis (Figure 5G-H). Together, these data provide *in situ* validation for scRNA-Seq-revealed metabolic heterogeneity within the basal compartment, and show basal cells in the wound neo-epidermis to be enriched for an OxPhos^low^ and glycolysis^high^ state compared to their UW counterparts.

### Pseudotemporal trajectory and RNA velocity analyses reveal basal state transition dynamics and wound-induced cellular plasticity

To pseudotemporally order the distinct basal cell states in the context of epidermal differentiation, we first performed Monocle 2 for trajectory analysis (Qiu et al., 2017; Trapnell et al., 2014) on all interfollicular epidermal cells, which include proliferating and non-proliferating basal cells as well as spinous cells (Figure S11). In both UW and WO skin, we observed three paths that extend from the *Col17a1*^Hi^ state: proliferating basal cells, GA state (transitioning through ER state), and spinous cells (Figure S11). *Col17a1*^Hi^ cells contribute in part to each of the observed paths (Figure S11A and S11C).

Since Monocle 2 is unable to determine the origin of the trajectory without prior knowledge, we also performed lineage predictions using our custom pipeline. First, we utilized uniform manifold approximation and projection (UMAP) for dimensional reduction (McInnes et al., 2018), where we visualized the cells by sample identity and corrected batch effects (Figure S12A-B, S12D, and 12DE; and see Methods section). This methodology when applied to a previously reported interfollicular epidermal 536-cell dataset revealed what appears to be a single-path trajectory just as reported (Joost et al., 2016) (Figure S12F). However, in our UW dataset, we observed 3 distinct and largely separated clusters composed of basal, proliferative basal, and spinous cells (Figure 6A-B). In the WO dataset, we identified the same 3 distinct epidermal clusters, but noted bridges between the basal and spinous clusters with overall less dramatic basal-spinous separation (Figure 6C-D).

**Figure 6:**
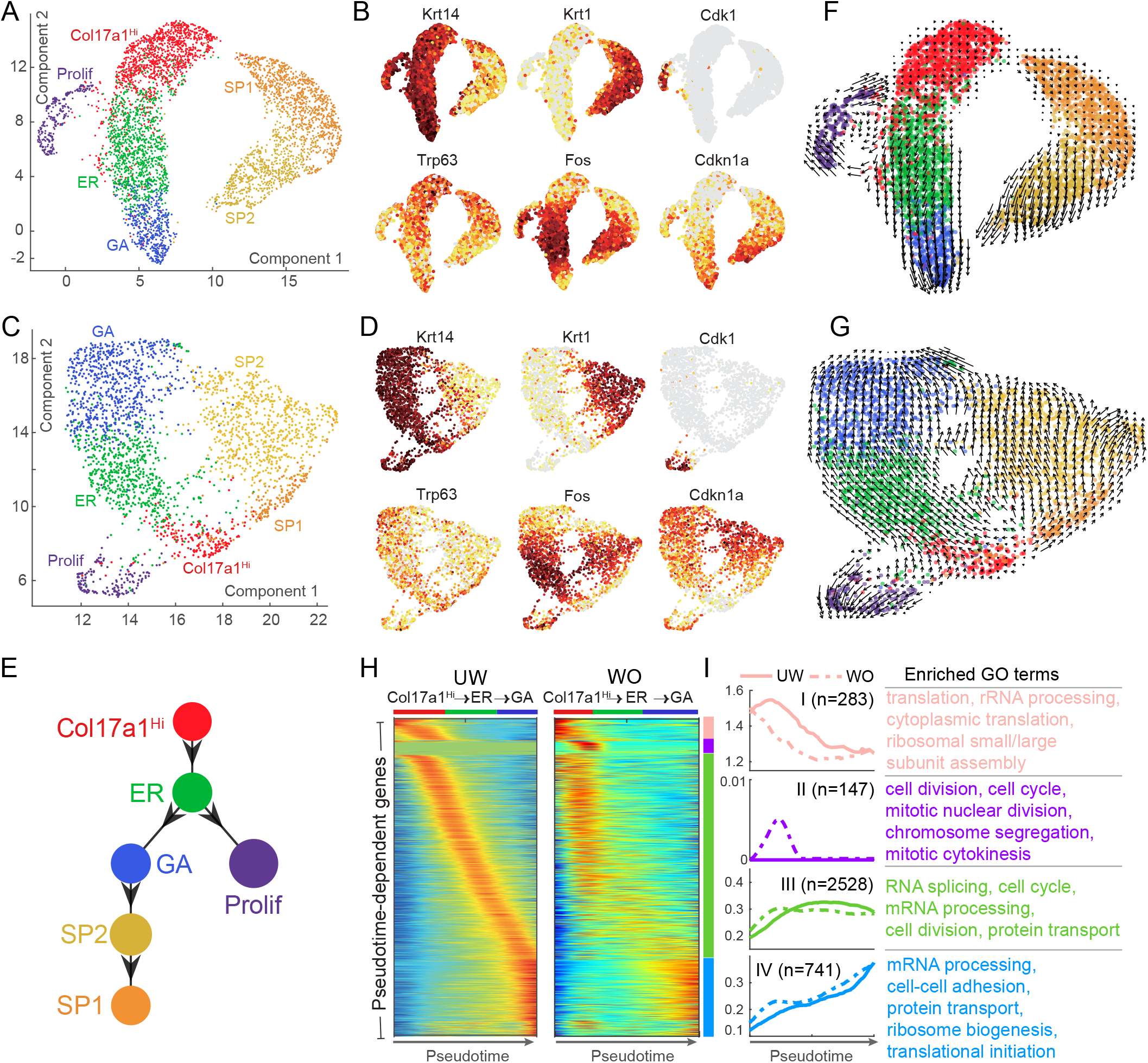
Lineage analysis of epidermal cells in UW and WO skin. A. UMAP dimensional reduction of all epidermal cells (non-proliferative basal, proliferative basal, and spinous) from UW sample. Cells are colored by the annotated identity. B. Feature plots of (A) for the indicated genes. Cells are colored by the normalized expression with dark red indicating highest expression. C. UMAP dimensional reduction of all epidermal cells (non-proliferative basal, proliferative basal, and spinous) from WO sample. Cells are colored by the annotated identity. D. Feature plots of (C) for the indicated genes. Cells are colored by the normalized expression with dark red indicating highest expression. E. scEpath-predicted lineage differentiation diagram using robust parameters. F. Projection of non-linear RNA velocity fields onto the UMAP space in (A). G. Projection of non-linear RNA velocity fields onto the UMAP space in (C). H. Pseudotemporal dynamics of the 3,699 UW pseudotime-dependent genes along the *Col17a1*^Hi^ to GA path in UW and WO samples. Cell identity (*Col17a1*^Hi^, ER or GA) was indicated on the top of each heatmap generated by the smoothed, normalized gene expression (for the colormap, blue and red colors indicate low and high expression, respectively.). I. Each row/gene was normalized to its peak value along the pseudotime. Distinct stages during pseudotime are represented by pink, purple, green, and blue bars. The average expression of each gene subcluster was shown for UW (solid lines) and WO (dashed lines) samples, respectively. The number of genes in each gene subcluster was as indicated, and the enriched GO terms in these different subclusters are listed.

Next, we used single cell Energy path (scEpath) to infer cell lineage relationships (see Methods section) (Jin et al., 2018). Single cell Energies (scEnergy) were calculated, projected onto UMAP space, and used to infer cell state transition probabilities and lineage relationships. We found that lower energies, while typically associated with committed/differentiated cell states (Jin et al., 2018; Teschendorff and Enver, 2017), are also associated with a quiescent cell state because the known quiescent Bu-HFSC (*CD34*^+^) population showed the lowest scEnergy of all the skin epithelial cell types (Figure 3D, 4E, and S4H; see Methods section). Using increasingly robust parameters, which incorporate increasing numbers of UW epidermal cells from each cell state to infer lineage progression, scEpath predicted a near-linear path that originates from the *Col17a1*^Hi^ basal cell state that displayed the lowest energy of all interfollicular epidermal cells (Figure 6E, S12C, and S12G). The *Col17a1*^Hi^ state transitions to the ER state, to the GA state, and then to the spinous cell populations (SP2 to SP1), whereas proliferative basal cells follow a side path that originates from the ER state (Figure 6E).

To further analyze the epidermal differentiation dynamics in UW skin, we performed RNA velocity analysis, a computational tool that can predict potential directionality and speed of cell state transitions based on levels of spliced and unspliced mRNA (La Manno et al., 2018; Svensson and Pachter, 2018) (see Method section). These transition dynamics are visualized by vectors (arrows), the direction of which relates to the direction of state transitions, and the length of which relates to the extent of change in RNA dynamics (La Manno et al., 2018; Svensson and Pachter, 2018). The *Col17a1*^Hi^ and SP1 states showed small RNA velocities (short or no arrows), which are known to associate with both quiescent and terminally differentiated cells (Svensson and Pachter, 2018; Zywitza et al., 2018), whereas the ER, GA, and SP2 states exhibited large RNA velocities (Figure 6F). Transition from the *Col17a1*^Hi^ state to the ER state represented a rapid activation in RNA dynamics (long arrows) (Svensson and Pachter, 2018) (Figure 6F). Moreover, the basal-to-spinous transition (Figure 6E) was characterized by dynamic forward and backward movements of the vectors between GA and SP2 states, which suggests uncommitted behaviors (Guo et al., 2018) before fully committing to a spinous fate (Figure 6F). Remarkably, the proliferative basal cells followed a cyclical trajectory, which originates from the border between *Col17a1*^Hi^ and ER states, and then returned to the *Col17a1*^Hi^ state (Figure 6F). These data suggest that epidermal basal cells normally transition through 3 distinct states before embarking on spinous differentiation, and this differentiation trajectory is fueled by the active proliferation of basal cells at the junction between *Col17a1*^Hi^ and ER states.

The cyclical dynamics of the proliferative UW basal cells was faithfully recapitulated in the WO sample (Figure 6G). While some of the WO *Col17a1*^Hi^ basal cells displayed slow RNA dynamics, an appreciable fraction of them and all of the SP1 cells exhibited apparently rapid RNA dynamics compared to their UW counterparts (Figure 6G). RNA velocity also provided evidence for enhanced cell-type fluidity in the WO sample, with bidirectional transitions between the basal and spinous cells detected at multiple cellular states that were not seen in the UW sample (Figure 6D and 6G). Overall, these data suggest that WO skin epidermal cells not only are generally more active but also exhibit increased plasticity and relaxed cell differentiation constraints compared to their UW counterparts.

We next sought to identify key molecular changes that may be important for pseudotemporal progression through basal cell states. scEpath identified 3699 and 3129 pseudotime-dependent genes (including basal state hallmark genes *Trp63*, *Fos*, and *Cdkn1a*) from the UW and WO dataset, respectively, that changed significantly as they transitioned through the basal cell states (Figure 6H, S13A, and S13B). Genes related to protein translation, rRNA processing, cell cycle, and cell division define the *Col17a1*^Hi^ to ER transition (Group I and II, Figure 6H and 6I). Moreover, genes related to mRNA processing, cell cycle, and protein transport define the *Col17a1*^Hi^/ER to GA transition (Group III), whereas those related to mRNA processing, cell adhesion, protein transport, and translation define the GA state (Group IV) (Figure 6H and 6I). While gene expression changes were sequential and gradual during basal state transitions in the UW sample, they appeared significantly earlier, were more abrupt and sometimes sporadic in the WO sample (Figure 6H, 6I, S13B, and S13C). Group II and III genes displayed the most dramatic differences between UW and WO datasets (Figure 6H and 6I), whereas TFs, which represent less than 1% of the total 3699 pseudotime-dependent genes, showed little difference (Figure S13D). Thus, cell cycle and post-transcriptional events underlie the wound-induced remodeling of basal cell state transitions.

The overall picture emerging from our computational lineage analysis is that epidermal basal cells in a primitive, *Col17a1*^Hi^ state first become “activated” (an early response-like state), at which point they can either enter active cell cycle and expand as progenitor cells, or undergo growth arrest and subsequently differentiate into spinous cells. While this multi-step basal-spinous differentiation trajectory is largely maintained during wound healing, cell fate plasticity and differentiation fluidity are enhanced such that bidirectional conversions between basal and spinous cells are enabled.

## DISCUSSION

Epidermal stem/progenitor cells in the basal layer of mouse adult skin maintain epidermal homeostasis by constantly replacing differentiated cells (Gonzales and Fuchs, 2017). Basal cells also must be able to alter their proliferative and migratory dynamics in the context of wound healing to facilitate re-epithelialization (Haensel and Dai, 2018). In this study, we have combined scRNA-Seq with *in situ* and *ex vivo* imaging methods to uncover previously unknown molecular and metabolic heterogeneities of the epidermal basal cells in normal skin and during the re-epithelization stage of wound healing. We have identified four distinct basal cell states, namely *Col17a1*^Hi^ (relatively quiescent), ER (early response), proliferative, and GA (growth arrested), in mouse back skin that shift gene expression, relative proportions, and spatial location during wound healing. Using multiple computational pipelines, we order these distinct states into a sequential differentiation hierarchy, revealing cyclical transitions of non-growth-arrested basal cells through the proliferative state, as well as faster cell state dynamics and greater differentiation plasticity during wound healing.

To date, several published studies have used scRNA-Seq to characterize the unknown heterogeneities within mammalian skin epithelia. These studies have provided a general molecular and cellular categorization of the various epithelial components of the mouse (Joost et al., 2016, 2018) and human (Cheng et al., 2018) skin, generated key insights into how the fates of a small number of epidermal or HF stem and progenitor cells evolve during normal regeneration and wound healing (Joost et al., 2018; Yang et al., 2017), and unearthed p63-regulated molecular and cellular events in the developing mouse epidermis (Fan et al., 2018). Our work adds to this growing list by providing the first comprehensive study of the transcriptional and metabolic heterogeneities of interfollicular epidermal basal cells in both normal and wounded skin.

Our discovery of a *Col17a1*^Hi^ basal cell state is thought-provoking. This transcriptional state is associated with high quiescence, high oxidative phosphorylation, but low EMT, low differentiation, low hypoxia/inflammation, low scEnergy, and low RNA dynamics. These molecular characteristics are suggestive of a relatively quiescent, primitive stem/progenitor cell state, a notion supported by its placement as the initial state in the epidermal lineage (this study), and its high expression of *Col17a1*, which encodes a marker of long-term epidermal stem cells that can outcompete other cells (Liu et al., 2019). The *Col17a1*^Hi^ state is enriched for genes such as *Trp63*, a master regulator of various aspects of epidermal development, including the initial specification from simple epithelia, promotion of stratification, proliferation, as well as terminal differentiation (Li et al., 2019; Mills et al., 1999; Pattison et al., 2018; Truong et al., 2006; Yang et al., 1999). The reduction of basal cells in the *Col17a1*^Hi^ state during wound re-epithelization implicates their mobilization in the case of increased demands for cellular outputs.

ER genes such as *Fos* and *Jun* are known as stress response genes that can be upregulated by flow cytometry (van den Brink et al., 2017). A transient upregulation of such “immediate early” genes during embryonic and adult wound healing has been previously suggested to represent a “kick-start” mechanism to initiate the repair process (Grose et al., 2002). We were able to detect epidermal basal cells *in situ* that express ER-associated Fos protein or *Id1* mRNA, indicating that an ER basal cell transcriptional state likely exists even in the intact, unwounded tissue. In all gene expression and lineage prediction analyses, these cells occupy an intermediate position between *Col17a1*^Hi^ and GA states, raising the possibility that ER is an obligatory transition state when dormant cells become activated to proliferate or migrate.

Basal cells that have committed to differentiation (expressing early differentiation marker *Involucrin* or *Inv*) have been identified via lineage tracing (Mascré et al., 2012). Although *Inv* expression was not detectable in our scRNA-seq-identified GA basal cells, it is likely that the GA state in normal epidermis is post-mitotic, most ready to commit to differentiation, and most prone to migrate upward compared to the other basal cell states. The high expression of EMT, glycolysis, hypoxia, and inflammation genes associated with the GA state implicates it as being the most ready to respond to extracellular signals from the ever-changing tissue microenvironment. Indeed, the relative number of GA-state basal cells is significantly increased upon wound healing and GA cells preferentially localize to the wound migrating front that is closest to the hypoxic wound bed and the infiltrating immune cells, as suggested by our scRNA-seq and RNAscope data. As such, our results is in keeping with the previous finding made using intravital imaging that migration and proliferation are spatially separated in the healing wound, with migrating cells at the tips of the growing neo-epidermis being generally devoid of proliferative activity (Park et al., 2017). More importantly, our data suggest that the wound repair process capitalizes on the existing transcriptional heterogeneity of the normal epidermal basal cells but redirects it towards a spatially coordinated program of proliferation, migration, and metabolism to facilitate efficient re-epithelialization.

We employed several computational tools to investigate basal cell dynamics in epidermal lineage differentiation. In all cases, a sequential progression of basal cells through the *Col17a1*^Hi^, ER, and then GA states was predicted. The placement of GA as the transitional state between basal and spinous fates is consistent with ample experimental findings supporting the notion that basal cells undergo cell cycle exit when they terminally differentiate (Fuchs, 2008). The “back and forth” RNA velocity arrows between GA and SP2 cells raise the possibility that the basal-spinous transition can be reversible and dynamic (Guo et al., 2018) until a relatively stable spinous fate (SP1) is reached. Furthermore, significantly enhanced cell fate fluidity occurs in the wounded skin, evident by the overall less distinct gene expression differences between basal and suprabasal fates as well as multiple RNA velocity vector paths that bridge the different states of the basal and spinous cell clusters. Cellular plasticity during wound healing is well-documented, as cells of the hair follicle and sebaceous gland lineages can be reprogramed to gain an interfollicular epidermal fate (Park et al., 2017; Rognoni and Watt, 2018). Our findings highlight another layer of cell fate plasticity, namely bidirectional fluidity between basal and spinous fates at multiple transcriptional states within the interfollicular epidermal compartment.

Of great interest is our identification of a distinct pool of proliferating basal cells as a separate path that forms a loop with, and thus fueling, the rest of the basal cells. The unique location and behavior of these cells in the lineage trajectories, namely at the border and able to rapidly and reversibly cycle between *Col17a1*^Hi^ and ER basal cell states, suggests that 1) *Col17a1*^Hi^ and ER cells are predominantly responsible for generating more basal cells, whereas GA cells as a bulk population likely have reached a point of no return such that they can no longer re-enter the cell cycle to serve as a major source of basal cell self-renewal; 2) passage through the *Col17a1*^Hi^-ER border might be critical for the active proliferation of the otherwise slow cycling adult epidermal basal cells; and 3) basal stem/progenitor cells can readily switch back and forth between quiescence and activity during adult epidermal homeostasis.

Collectively, our findings reveal important new insights into the cellular and molecular dynamics of normal and wounded skin. The sequential progression of basal cells through three non-proliferative states, two of which are capable of reversibly entering active proliferation, is consistent with a revised “hierarchical-lineage” model of epidermal homeostasis that encompasses multiple possible stem/progenitor cell states. Our study lays the foundation for future spatial transcriptomics, high-resolution lineage tracing, and multiscale mathematic modeling to investigate how multi-state basal-spinous differentiation and the underlying regulatory/signaling networks contribute to the performance objective of adult epidermis, namely maintaining a homeostatic, functional epidermis that can robustly regenerate itself upon injury – a goal that is difficult to attain under pathological conditions.

## METHODS

### Mice

*K14-Cre* transgenic mice have been previously described (Andl et al., 2004). *ROSA^mTmG^* and C57BL/6J mice are from the Jackson Laboratory (Stock #s 007576 and 000664, respectively). Only female mice were used in this study. All experiments have been approved and abide by regulatory guidelines of the International Animal Care and Use Committee (IACUC) of the University of California, Irvine.

### Wounding

For single cell experiments, mice were anesthetized using isoflurane (Primal Healthcare; NDC-66794-017-25), backs shaved, and then a 6-mm punch (Integra; 33-36) was used to generate a full-thickness wound on each side of the mouse. For FLIM-based wounding experiments, mice were anesthetized using isoflurane, backs shaved, and Nair was applied to the backs of the shaved mice for complete hair removal. A 6-mm punch was used to generate a full-thickness wound on each side of the mouse. Four days after wounding, wound samples containing the wound and surrounding un-wounded skin regions were excised for analysis.

### Single cell isolation for scRNA-Seq

For UW back skin, mice were shaved, back skin removed, fat scrapped off, and then skin was minced into pieces less than 1 mm in diameter. For WO back skin, skin was removed, large pieces of fat attached to underside of wound were carefully removed, a 10-mm punch (Acuderm; 0413) was then used to capture the wound and a portion of un-wounded skin adjacent to the wound. The wounds were then minced into pieces less than 1 mm in diameter. The minced samples were placed in 15-mL conical tubes and digested with 10 mL of collagenase mix [0.25% collagenase (Sigma; C9091), 0.01M HEPES (Fisher; BP310), 0.001M Sodium Pyruvate (Fisher; BP356), and 0.1 mg/mL DNase (Sigma; DN25)]. Samples were incubated at 37 °C for 2 hours with rotation, and then filtered with 70-μm and 40- μm filters, spun down, and resuspended in 2% FBS. Cells were stained with SytoxBlue (Thermo Fisher; S34857) as per manufacturer’s instructions and live cells (SytoxBlue-negative) were sorted using BD FACSAria Fusion Sorter.

### Single cell library generation

FACS-sorted cells were washed in PBS containing 0.04% BSA and resuspended to a concentration of approximately 1,000 cell/µL. Library generation was performed following the Chromium Single Cell 3ʹ Reagents Kits v2 User Guide: CG00052 Rev B. where we target 10,000 cells per sample for capture. Each library was sequenced on the Illumina HiSeq4000 platform to achieve an average of approximately 50,000 reads per cell.

### Processing and clustering analysis of scRNA-seq data

FASTQ files were aligned utilizing 10x Genomics Cell Ranger 2.1.0. Each library was aligned to an indexed mm10 genome using Cell Ranger Count. Cell Ranger Aggr function was used to normalize the number of mapped reads per cells across the libraries. Clustering of cells was performed using the Seurat R package (Satija et al., 2015). Briefly, initial bulk samples containing single cell data matrices were column-normalized and log-transformed. Quality control parameters were used to filter cells with 200-5000 genes with a mitochondrial percentage under 10%. Replicates for UW and WO samples were merged and then corrected using the MultiCCA function. To identify cell clusters, principle component analysis (PCA) was first performed and the top 10 PCs with a resolution = 0.6 were used to obtaining 15 and 14 clusters for the UW and WO samples, respectively. For subclustering of epithelial cells, we first identified epithelial clusters from UW or WO replicate using the top 10 PCs with resolution = 0.6 and then subset out the appropriate epithelial clusters. Replicates of these epithelial clusters were then merged using MultiCCA function again using 10 PCs with resolution = 0.6. For subclustering of basal cells, we performed batch correction using the Bayesian-based method ComBat from the sva R package. The corrected data were used for further clustering analysis. Briefly, for the UW sample, the top 23 PCs were used for clustering and 3 subclusters were obtained with a resolution = 0.8. For the WO sample, the top 26 PCs were used and 3 subclusters were obtained with a resolution = 0.3. Marker genes were determined with p-value < 0.01 and log(fold-change) > 0.25 by performing differential gene expression analysis between the clusters using Wilcoxon rank sum test. To present high dimensional data in two-dimensional space, we performed t-SNE analysis using the results of PCA with significant PCs as input.

### Gene Scoring

For gene scoring, gene sets were acquired from the MSigDB database, and others (α5 integrin-expressing cell and quiescence scoring) were generated from published literature (Aragona et al., 2017; Cheung and Rando, 2013). The lists of genes in each gene set are listed in Table S8. The AddModuleScore function was then used to score various cell clusters.

### Random forest classifier

Using the Seurat R Package 2.2.0, we employed the ClassifyCells function, which relies on the Ranger package to build a random forest suited for high dimensional data. Training class was based on identities of the basal cells from the UW sample, which was subsequently applied to the basal cells from the WO sample.

### Pseudotime and trajectory analysis

We performed pseudotemporal ordering of all interfollicular epidermal cells, including proliferative and non-proliferative basal cells and spinous cells, using two methods: Monocle 2 (Qiu et al., 2017) and scEpath (Jin et al., 2018). For Monocle 2, batch effect information was passed into the residualModelFormulaStr option in the “reduceDimension” function. The scEpath method can quantify the energy landscape using scEnergy, which quantitatively measures the developmental potency of single cells (Jin et al., 2018) and was used in our analysis to predict the initial state in pseudotime. Pseudotemporal ordering was performed on batch corrected data, and batch correction was performed for both UW and WO samples using Combat. The corrected data was used as an input for dimension reduction using PCA and then UMAP. Based on this reduced UMAP space, scEpath infers lineage relationships between cell states via predicted transition probabilities and pseudotime (using a principal curve-based approach). To improve the robustness of estimating the centroid of cell clusters, scEpath defines metacell consisting of a set of cells that occupies θ1 percent of the total energy in each cluster. The centroid of the metacell in 2D was used to further infer the cell lineages. The inferred cell lineages of UW samples were robust to a higher portions of cells that were used for defining metacells. To investigate the global dynamic changes in gene expression, scEpath also identifies pseudotime-dependent genes by creating a smoothed version of gene expression using a cubic spline (Jin et al., 2018). To analyze pseudotime-dependent TFs, we used genes that are annotated in the Animal TF Database (AnimalTFDB 2.0) (Zhang et al., 2015).

### RNA velocity analysis

RNA velocity was calculated based on the spliced and unspliced counts as previously reported (La Manno et al., 2018), and cells that were present in the pseudotemporal ordering were used for the analysis. We used the R implementation “velocyto” with a modified dynamical model to perform RNA velocity analysis. *La Manno et al*. used a linear model to relate abundance of pre-mRNA U(t) with abundance of mature mRNA S(t):

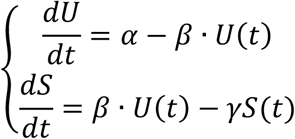

In this model, mRNA abundance over time (represented as dS/dt) is the velocity of gene expression. Given that the molecular regulatory mechanisms between pre-mRNA and mature mRNA are complicated, and in many molecular networks more commonly we observe non-linear (e.g. switch-like) responses, we proposed a nonlinear model of RNA velocity for the effects of pre-mRNA on the abundance of mature mRNA based on Michaelis–Menten kinetics. The nonlinear RNA velocity model is formulated as:

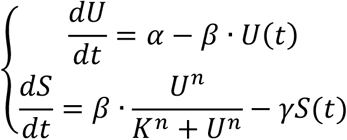

where n is the Hill coefficient (describing cooperativity) and K is a constant. We set n and K to be 1 and 0.5 in all the analyses below.

In both linear and nonlinear analysis, RNA velocity was estimated using gene-relative model with k-nearest neighbor cell pooling (k = 30). Velocity fields were then projected onto a low dimensional space (e.g. UMAP or the low-dimensional space produced by Monocle 2). Parameter n-sight, which defines the size of the neighborhood used for projecting the velocity, was set to 500.

### FLIM and data analysis

Freshly excised skin was placed in a glass bottom microwell dish (MatTek Corporation; PG-35g-1.5-14-C) and imaging was performed using a 63X Oil 1.4NA lens (Zeiss) on a Zeiss LSM 880 microscope coupled to a Ti:Sapphire laser system (Spectra Physics, Santa Clara CA, USA, Mai Tai HP). External hybrid photomultiplier tubes (Becker & Hickl; HPM-100-40) and ISS A320 FastFLIM system (ISS, Urbana-Champaign, Illinois) were used for Phasor Fluorescence Lifetime Imaging Microscopy (Colyer et al., 2008; Digman et al., 2008; Stringari et al., 2015). A 690 nm internal dichroic filter (Zeiss) was used to separate the fluorescence emission from the laser excitation. The fluorescence emission was reflected onto a 495LP dichroic mirror and subsequently a 460/80 nm bandpass filter (Semrock; FF02-460/80-25) before the external detector to filter the NADH fluorescence emission. Images were acquired using unidirectional scan, 16.38 us pixel dwell time, 256 x 256 pixels per frame, and 58.67um field of view. All images were acquired within 1.5 hours of animal death.

The phasor plot method provides a fit-free, unbiased way of analyzing FLIM data quantitatively. FlimBox, developed by the Laboratory for Fluorescence Dynamics at UC Irvine, records the photon counts per pixel in a number of cross-correlation phase bins called the phase histogram used for the Digital Frequency Domain FLIM method. The phase histogram is processed by the fast Fourier transform to produce the phase delay ϕ and modulation ratio m of the emission relative to the excitation from which the G and S coordinates calculated at each pixel of the image are represented in the phasor plot.

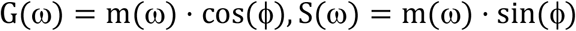

Data analysis was performed with Globals for Images (SIMFCS 4.0) software developed at the Laboratory for Fluorescence Dynamics. We used coumarin 6 (Sigma-Aldrich; 546283), with known lifetime of 2.5ns, for calibration of the instrument response function.

Quantification of the average NADH phasor per region of interest was calculated using the built-in masking feature in SimFCS 4.0. This masking feature averages the lifetime (τ) of all pixels included within a designated region of interest (ROI). SimFCS converts G and S coordinates of the phasor plot into the fraction of bound by calculating the distance of the ROI average τ to the theoretical lifetime τ of bound NADH (τ= 3.4 ns), divided by the total distance between free NADH (τ= ns) and bound NADH. An ROI within the boundary of each cell demarked by GFP expression was drawn to estimate the free/bound NADH ratio for each cell within a field of view for all images. The fraction bound values obtained from SimFCS 4.0 were then converted to free/bound ratio NADH for each ROI as a measure of metabolism based on previous work (Cinco et al., 2016; Kim et al., 2016; Mah et al., 2018; Stringari et al., 2012, 2015).

### Morphology and immunostaining

For histological analysis, mouse back skin was shaved, removed, fixed in 4% paraformaldehyde (MP; 150146) in 1X PBS, embedded in paraffin, sectioned, and stained with hematoxylin and eosin (H/E). For indirect immunofluorescence, mouse back skin was freshly frozen in OCT (Fisher; 4585), sectioned at 5 μm, and staining was performed using the appropriate antibodies and DAPI (Thermo Fisher; D1306: 1:1000). The following primary antibodies were used: Ki67 (Cell Signaling, D3B5, 1:1000), K14 (chicken, 1:1000; rabbit, 1:1000; gift of Julie Segre, National Institutes of Health, Bethesda), Fos (Santa Cruz Biotechnology, sc271243, 1:100).

### RNAScope, data analysis and presentation

For RNAScope, we utilized the Multiplex Fluorescent v2 system (ACD; 323100). Mouse back skin or wounds were freshly frozen in OCT (Fisher; 4585) and sectioned at 10 μm. Sections were fixed at room temperature for 1 hour with 4% paraformaldehyde (Electron Microscopy Sciences; 15715-S), which was diluted from stock with 1x DPBS (Corning Cellgro; 21-031-CM). After fixation, standard RNAScope protocols were used according to manufacturer’s instructions. The following probes were used: *Krt14* (ACD; 422521-C3), *Trp63* (ACD; 464591-C2), *Cdkn1a* (ACD; 408551-C1), and *Id1* (ACD; 312221-C3). We quantified the fluorescence intensity of the basal cells (stained positive for anti-K14 antibody and adjacent to the basement membrane or wound bed) sequentially in both UW and WO (from the wound margin to the tip of the migrating front) samples to obtain relative spatial information.

We used Gaussian process regression (GPR), a non-parametric method to fit observations, to visualize the major trends of data by controlling the smoothness of the model. GPR uses kernels to measure similarity between inputs based on their distances, and inputs with high similarity should have similar output from the fitted model. We used the implementation of GPR in scikit-learn package (Pedregosa et al., 2011; Rasmussen and Williams, 2006). The Matérn kernel is used for similarity measurement and a white noise kernel is included to accommodate noise in the data.

We used a commonly used index Moran’s I to analyze spatial autocorrelation (the similarity/dissimilarity of a variable in spatial neighbors). The local Moran’s I at location j is computed as:

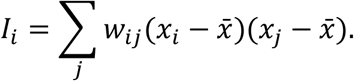

The global Moran’s I is simply the summation:

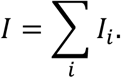

We computed the spatial autocorrelation for our sample with 2-nearest neighbors, in other words,

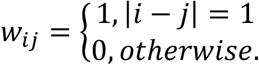

The sample is regarded as a series of cells aligned in one dimension and |*i* - *j*| = 1 means cell *i* and cell *j* are adjacent to each other.

Given a collection of values, BASC method (Hopfensitz et al., 2012) first sorts the values to obtain an initial step function representation. This step function is then iteratively refined until there are only two steps. It can be roughly understood as finding the strongest discontinuity point in data. The R implementation of this package “Binarize” is used with algorithm option B to determine thresholds for binarization of the markers.

## ABBREVIATIONS

Bulge hair follicle stem cells: (Bu-HFSCs)
Canonical Correlation Analysis: (CCA)
Dendritic cell: (DC)
Early response: (ER)
Epithelial-to-mesenchymal transition: (EMT)
Fluorescence lifetime imaging microscopy: (FLIM)
Gaussian process regression: (GPR)
Gene ontology: (GO)
Growth arrested: (GA)
Hair follicle: (HF)
Label-retaining cell: (LRC)
Single cell RNA-sequencing: (scRNA-Seq)
t-distributed stochastic neighbor embedding (tSNE) Transcription factor: (TF)
Two-photon excitation: (TPE)
Uniform manifold approximation and projection: (UMAP)
Un-wounded: (UW)
Wounded: (WO)

## ACKNOWLEDGEMENT

We thank the Genomics High Throughput Facility and the Institute for Immunology FACS Core Facility at the University of California, Irvine (UCI) for expert service. FLIM experiments reported in this publication were performed at the Laboratory for Fluorescence Dynamics, which is supported jointly by the NIGMS/NIH (P41GM103540) and UCI. This work was supported by NIH Grants R01-AR068074 (X.D.), R01GM123731 (X.D.), U01-AR073159 (MPIs: Q.N., M.P., and X.D.); NSF Grants DMS1763272, DMS1562176 and Simons Foundation Grant 594598 (Q.N.).

## AUTHOR CONTRIBUTIONS

XD and DH conceived the study and designed the experiments; XD directed the project; DH, RC, PS, and MD performed all wet-lab experiments; DH, SJ, QN(guyen), ZC, YG, and ALM performed and assisted with computational analysis; QN(ie) supervised the pseudotime and trajectory analysis; RC performed FLIM analysis under EG’s supervision; QN(guyen) assisted with sample processing and sequencing under KK’s supervision; DH and XD wrote the manuscript with inputs from all authors.

## COMPETING INTERESTS

The authors declare no competing interests.

## SUPPLEMENTARY FIGURE LEGENDS

**Supplementary Figure 1:**
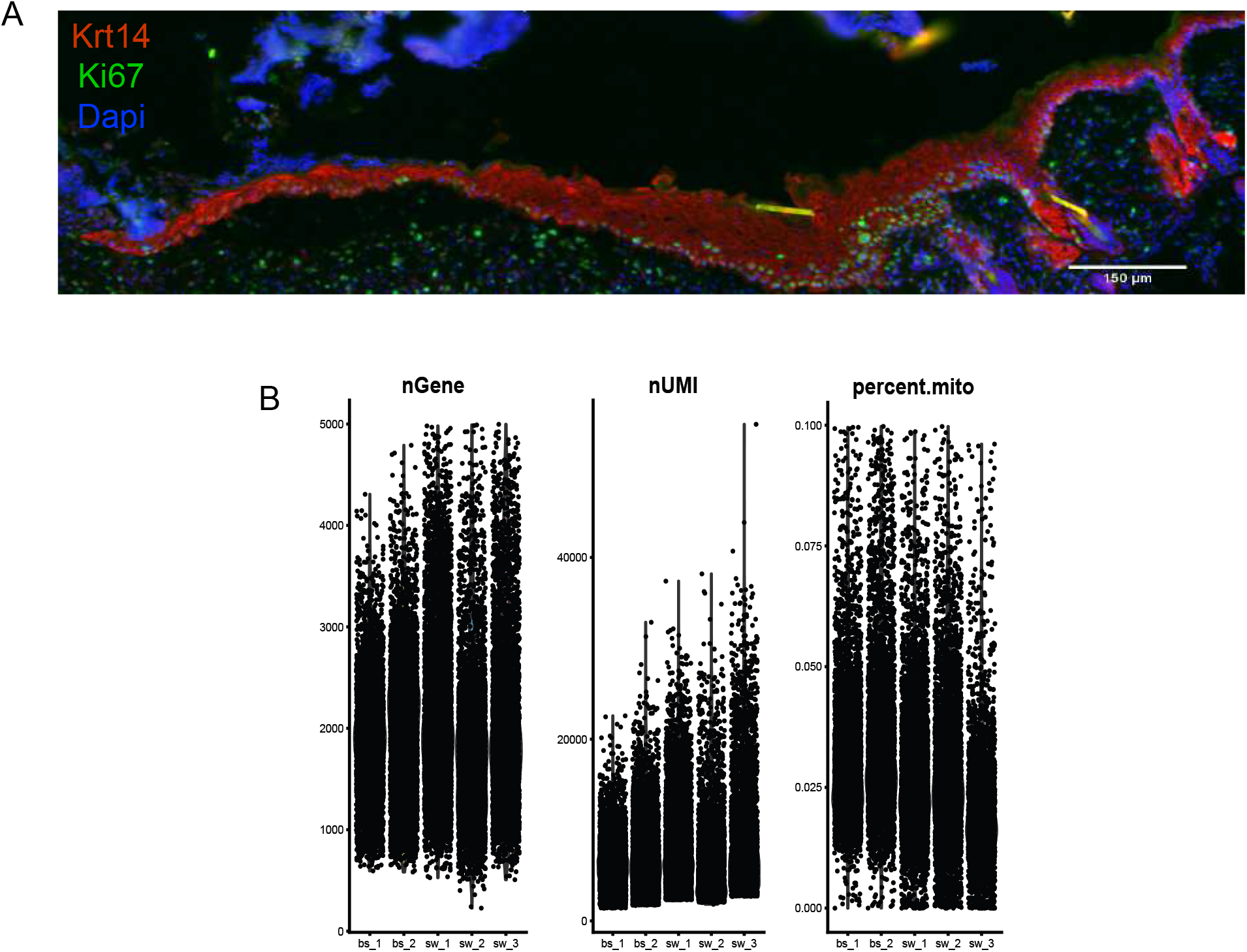
Proliferation dynamics and quality control metrics. A. Proliferation dynamics in equivalent sample taken for scRNA-Seq. Yellow dashed line indicates wound margin. B. Quality control metrics of UW and WO samples indicating the number of genes (nGene with 200-5000 range), number of UMIs (nUMI), and the percent mitochondrial genes (percent.mito under 10%).

**Supplementary Figure 2:**
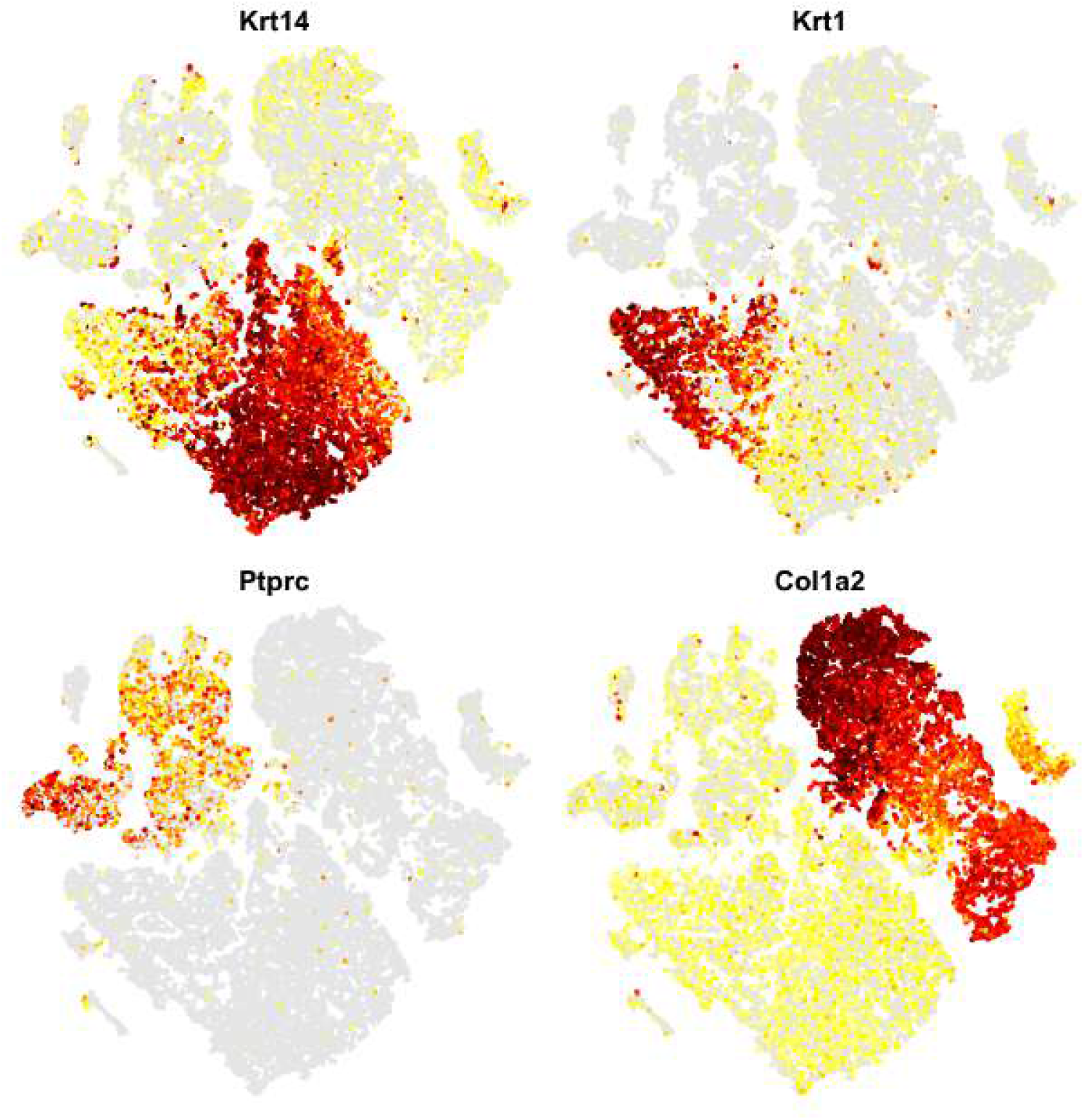
Feature plots of major cell type populations containing all samples (2 UW and 3 WO)

**Supplementary Figure 3:**
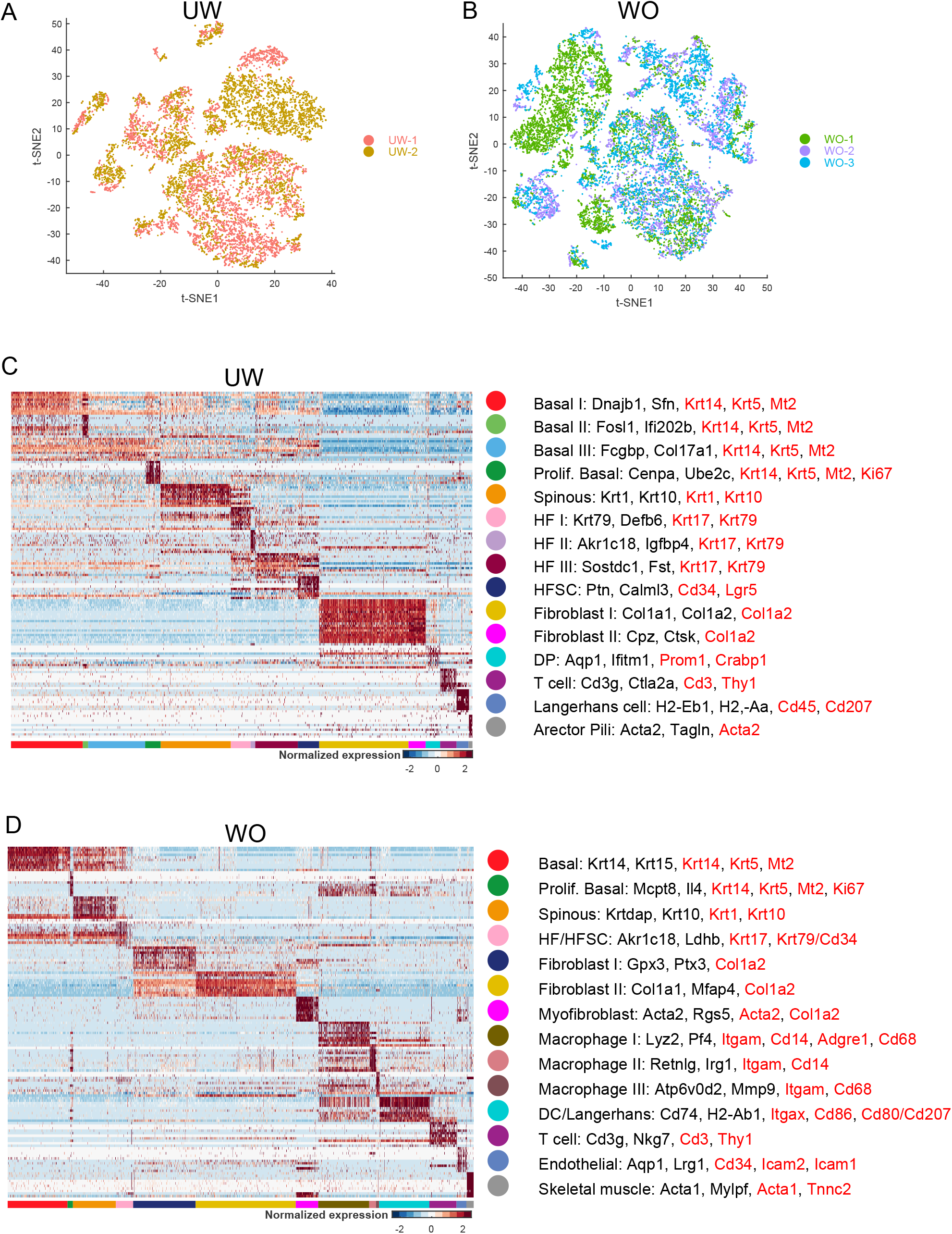
Replicate IDs and associated marker genes with heatmaps. A. tSNE plot containing 2 samples from un-wounded skin that were aggregated and then batch corrected using CCA. Colors correspond to each replicate sample identified by a unique color identifier. t-distributed stochastic neighbor embedding (tSNE) of cells from UW and WO samples showed that there was mixing between the replicate samples; however, unsurprisingly some residual batch effects still persisted after CCA correction due to individual variations B. tSNE plot containing 2 samples from un-wounded skin that were aggregated and then batch corrected using CCA. Colors correspond to each replicate sample identified by a unique color identifier. C. Heatmap for the top 10 genes enriched in each of the unique clusters from un-wounded skin. Genes listed in black represent the top 2 marker genes from each cluster where the genes listed in red represent the genes used in the final identification of the cluster cell type. D. Heatmap for the top 10 genes enriched in each of the unique clusters from wounded skin. Genes listed in black represent the top 2 marker genes from each cluster where the genes listed in red represent the genes used in the final identification of the cluster cell type.

**Supplementary Figure 4:**
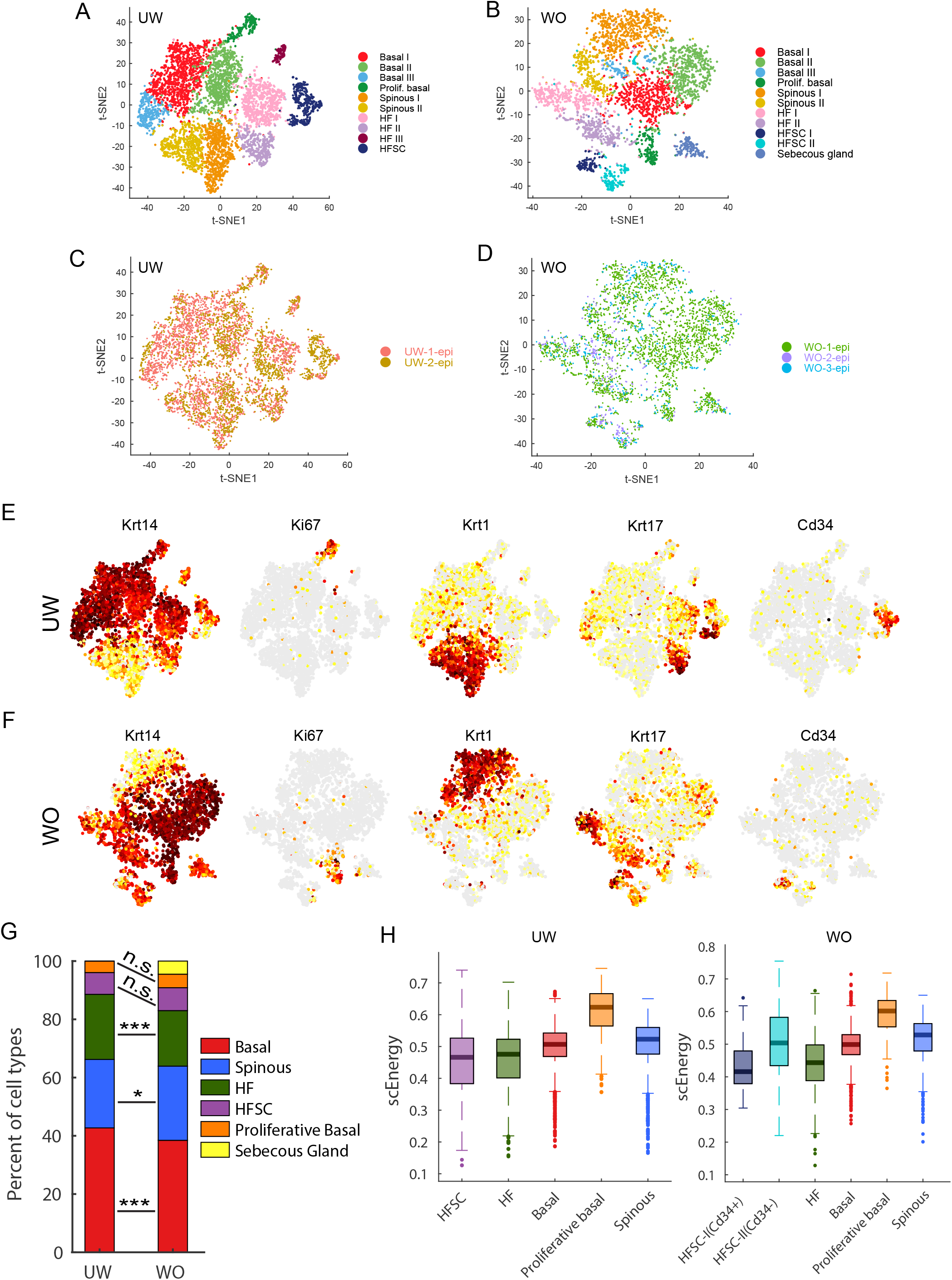
scRNA-Seq reveals minor changes in epithelial cellular makeup during wound healing. A. tSNE plot containing epithelial cells from 2 samples from un-wounded skin that were aggregated and then batch corrected using CCA. B. tSNE plot containing epithelial cells from 3 samples from wounded skin that were aggregated and then batch corrected using CCA. C. tSNE plot from (A) where colors correspond to each replicate sample identified by a unique color identifier. D. tSNE plot from (B) where colors correspond to each replicate sample identified by a unique color identifier. E. Feature plots highlighting genes from major epithelial cell types in UW sample. F. Feature plots highlighting genes from major epithelial cell types in WO sample. G. Bar graph representing the major epithelial cell type populations in the UW and WO samples. For statistical analysis, Chi-squared test was used (\**p* < 0.05; *** *p* < 0.0005). H. Boxplot indicating scEnergy values for different cell states from UW and WO samples containing epithelial cells from S4A and S4B, respectively (see Methods section). The quiescent Bu-HFSC population (CD34^+^) displayed lowest scEnergy values.

**Supplementary Figure 5:**
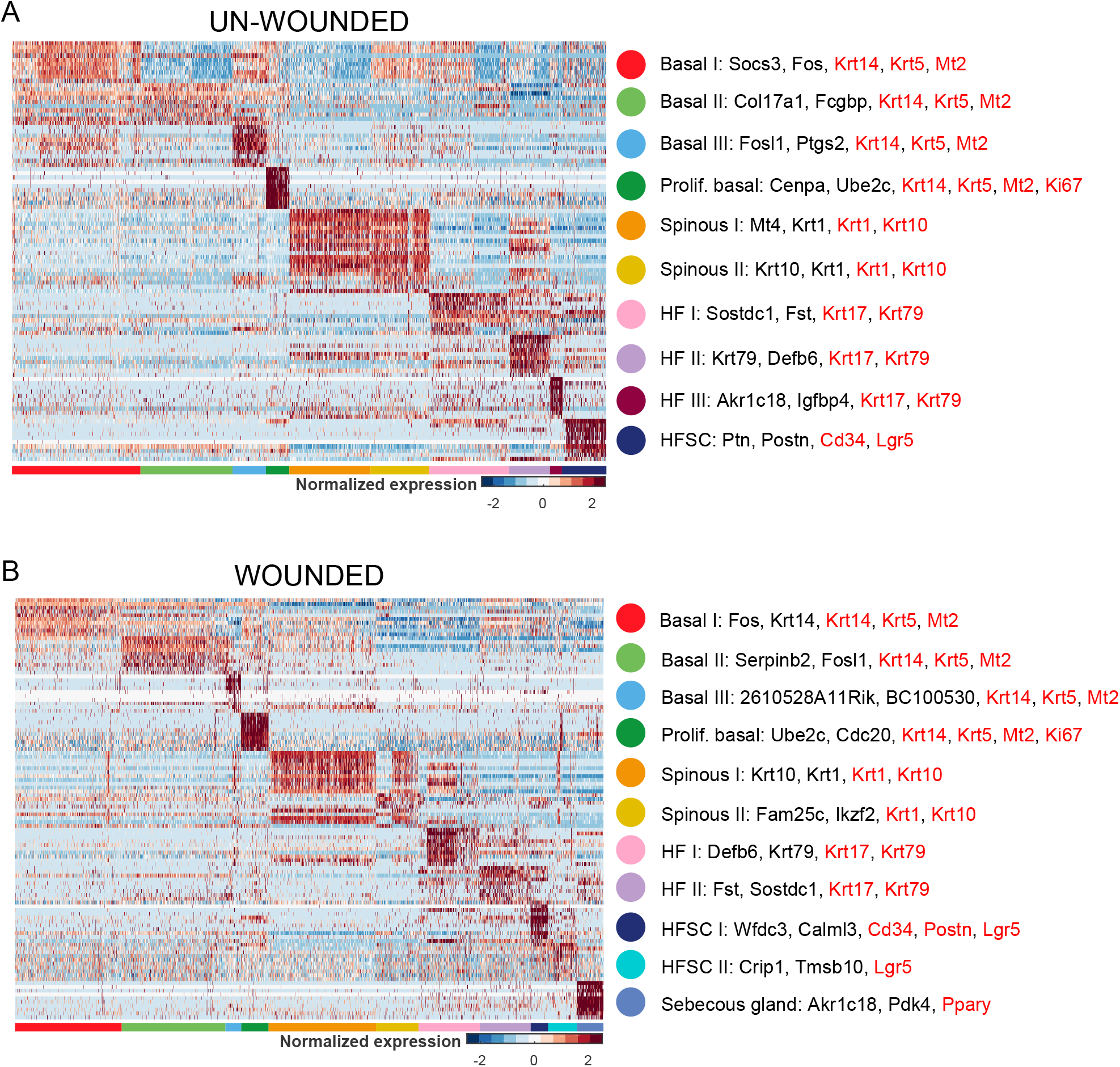
Associated marker genes with heatmaps for epithelial cells from UW and WO samples. A. Heatmap for the top 10 genes enriched in each of the unique epithelial clusters from un-wounded skin. Genes listed in black represent the top 2 marker genes from each cluster where the genes listed in red represent the genes used in the final identification of the cluster cell type. B. Heatmap for the top 10 genes enriched in each of the unique epithelial clusters from wounded skin. Genes listed in black represent the top 2 marker genes from each cluster where the genes listed in red represent the genes used in the final identification of the cluster cell type.

**Supplementary Figure 6:**
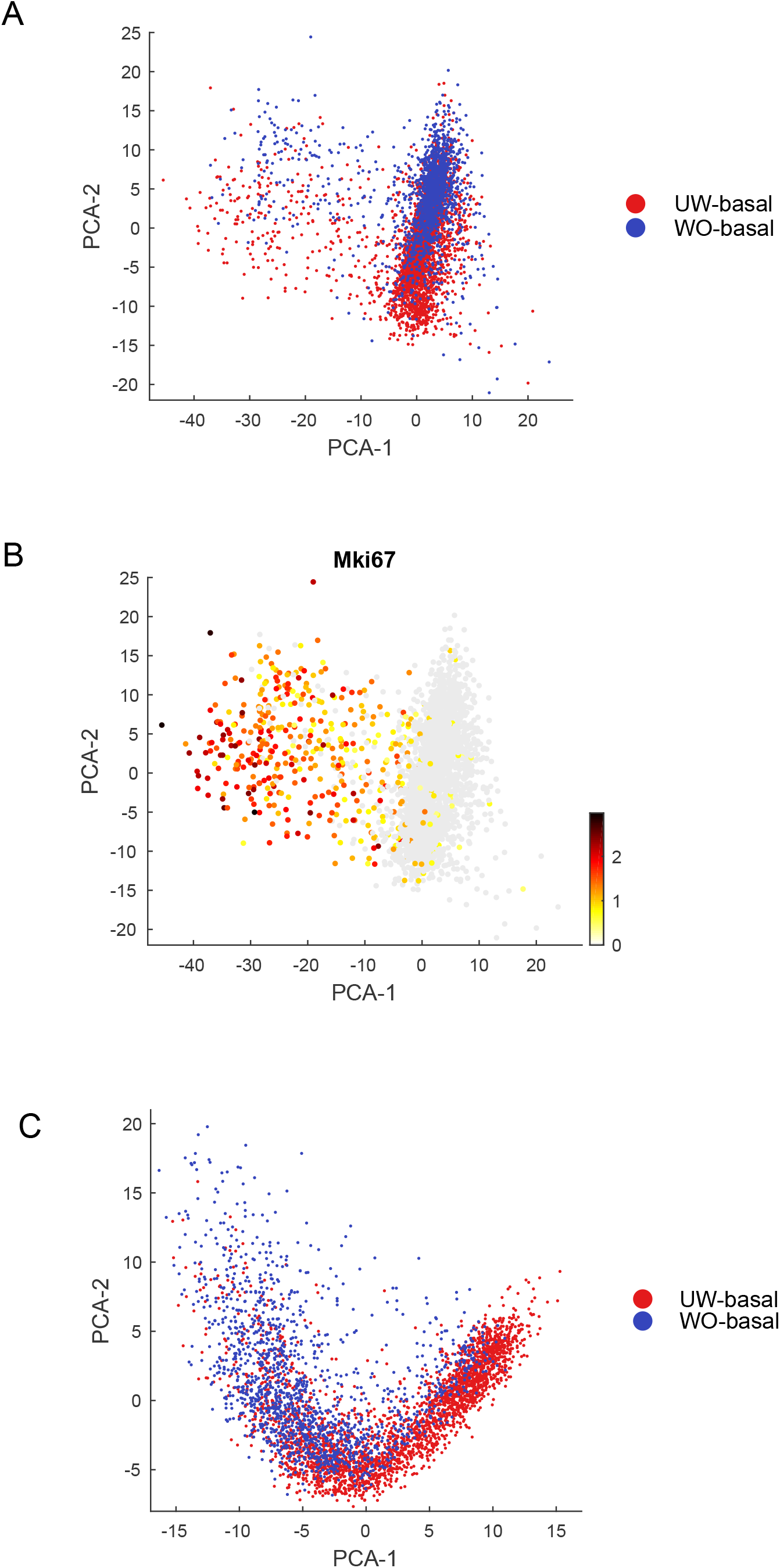
Proliferative basal cells have dramatically distinct gene expression patterns compared to non-proliferative basal cells. A. PCA analysis of total basal cells (proliferative and non-proliferative) from UW and WO samples. B. Feature plot utilizing PCA plot in (A) indicating the expression of proliferative marker *Mki67* in a subset of cells. C. PCA analysis of non-proliferative basal cells from UW and WO samples after removal of proliferative cells.

**Supplementary Figure 7:**
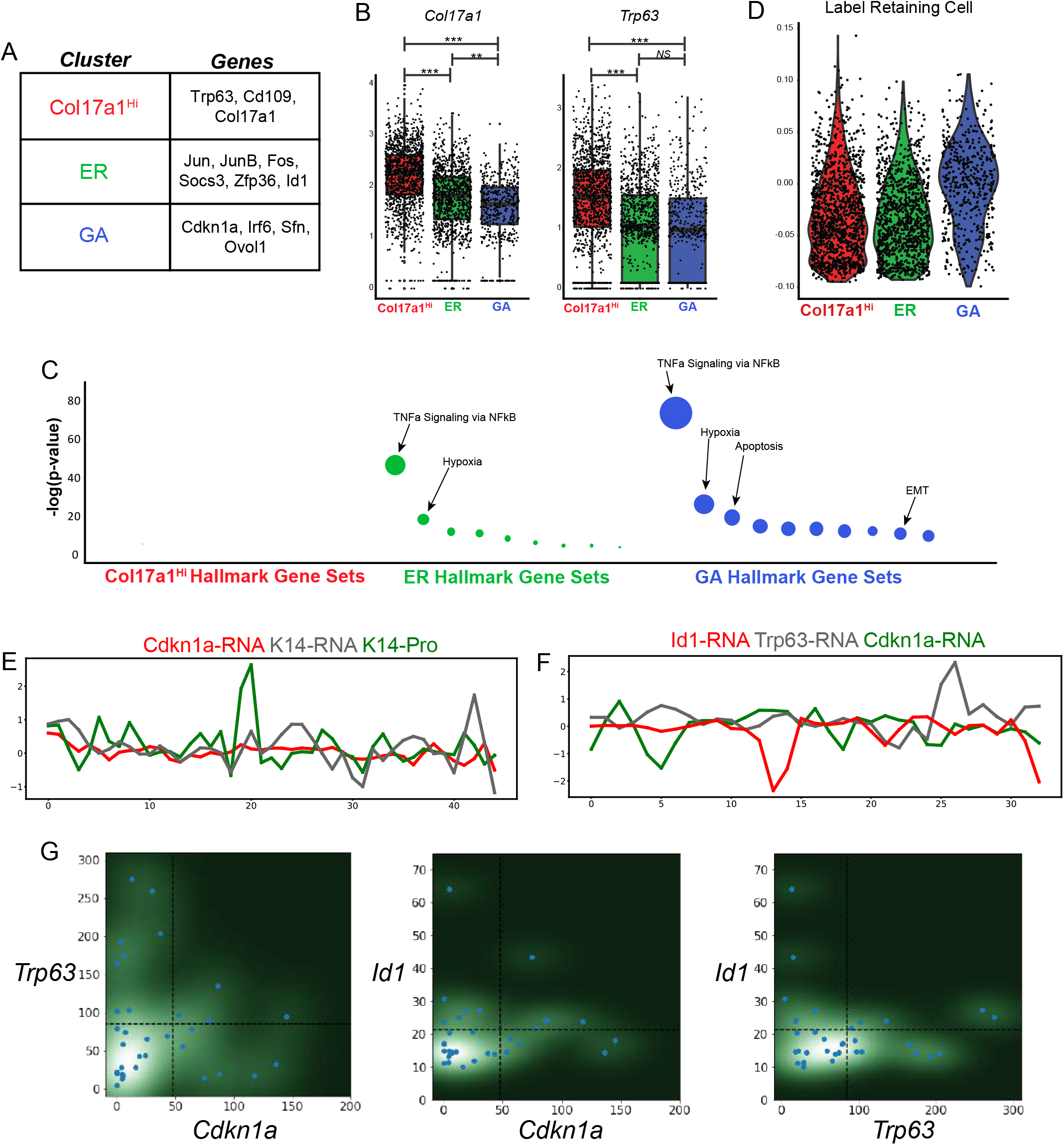
Basal cell analysis in UW skin. A. Key marker genes enriched in the *Col17a1*^Hi^, ER, and GA basal cell subcluster. B. Boxplots indicating mean value of *Col17a1* and *Trp63* expression in each basal cell subcluster. C. GO analysis of UW basal subclusters. D. LRC gene score for *Col17a1*^Hi^, ER, and GA basal cell subclusters. E. Local spatial autocorrelation analysis of RNAScope analysis of *Cdkn1a*, *Krt14*, and K14. F. Local spatial autocorrelation analysis of RNAScope analysis of *Cdkn1a*, *Trp63*, and *Id1*. G. Heatmap showing density estimation with dashed lines being cutoffs for binarization.

**Supplementary Figure 8:**
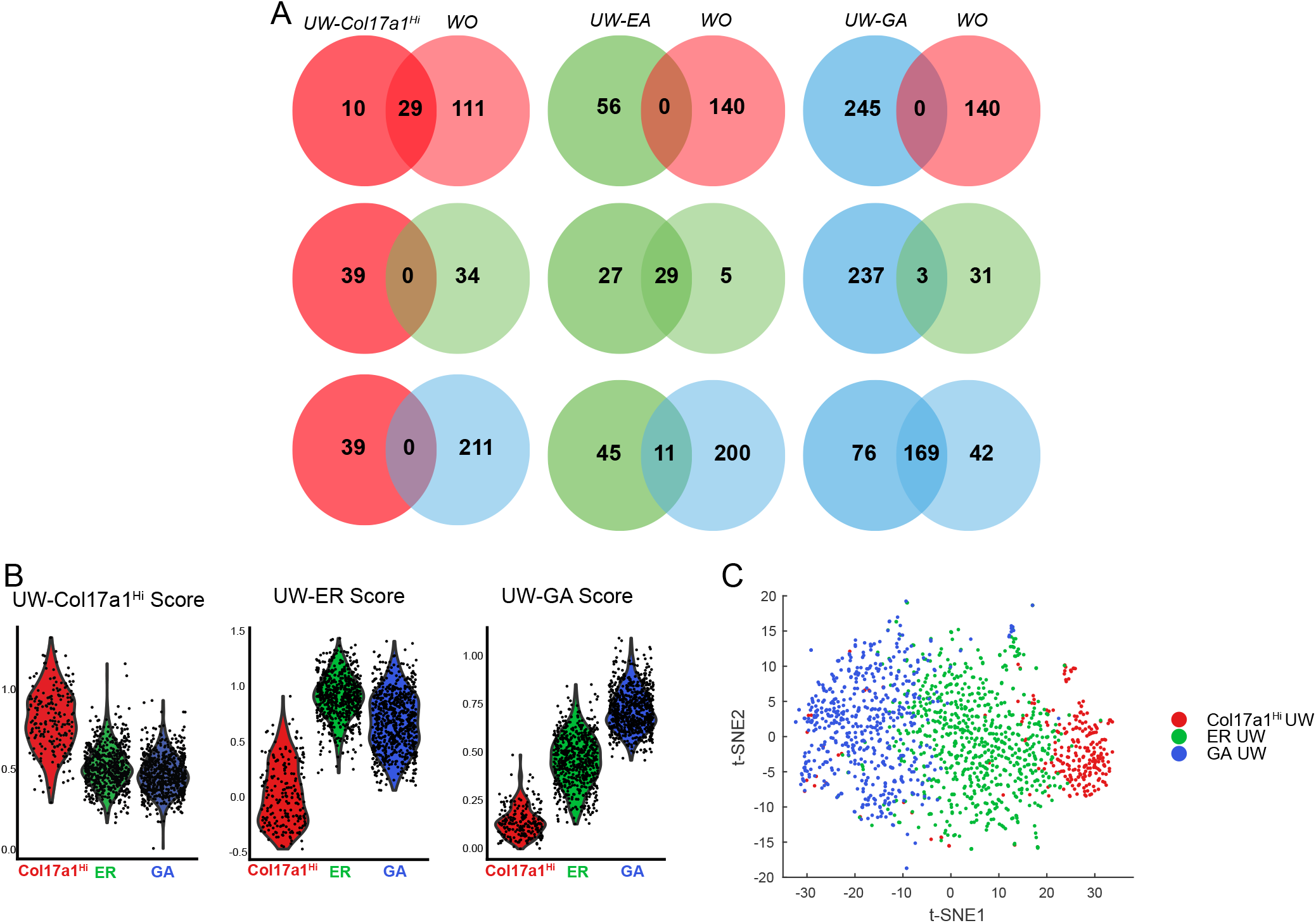
Basal cell states are similar in UW and WO samples. A. Marker gene overlap between the *Col17a1*^Hi^, ER, and GA basal cell subclusters from UW samples and basal cell subclusters from basal cell subclusters identified in WO sample. B. Scoring different basal cell subclusters from WO sample using marker genes from basal cell subclusters from UW samples. C. Random forest classification of basal cells from WO sample using basal cells from UW sample as training class.

**Supplementary Figure 9:**
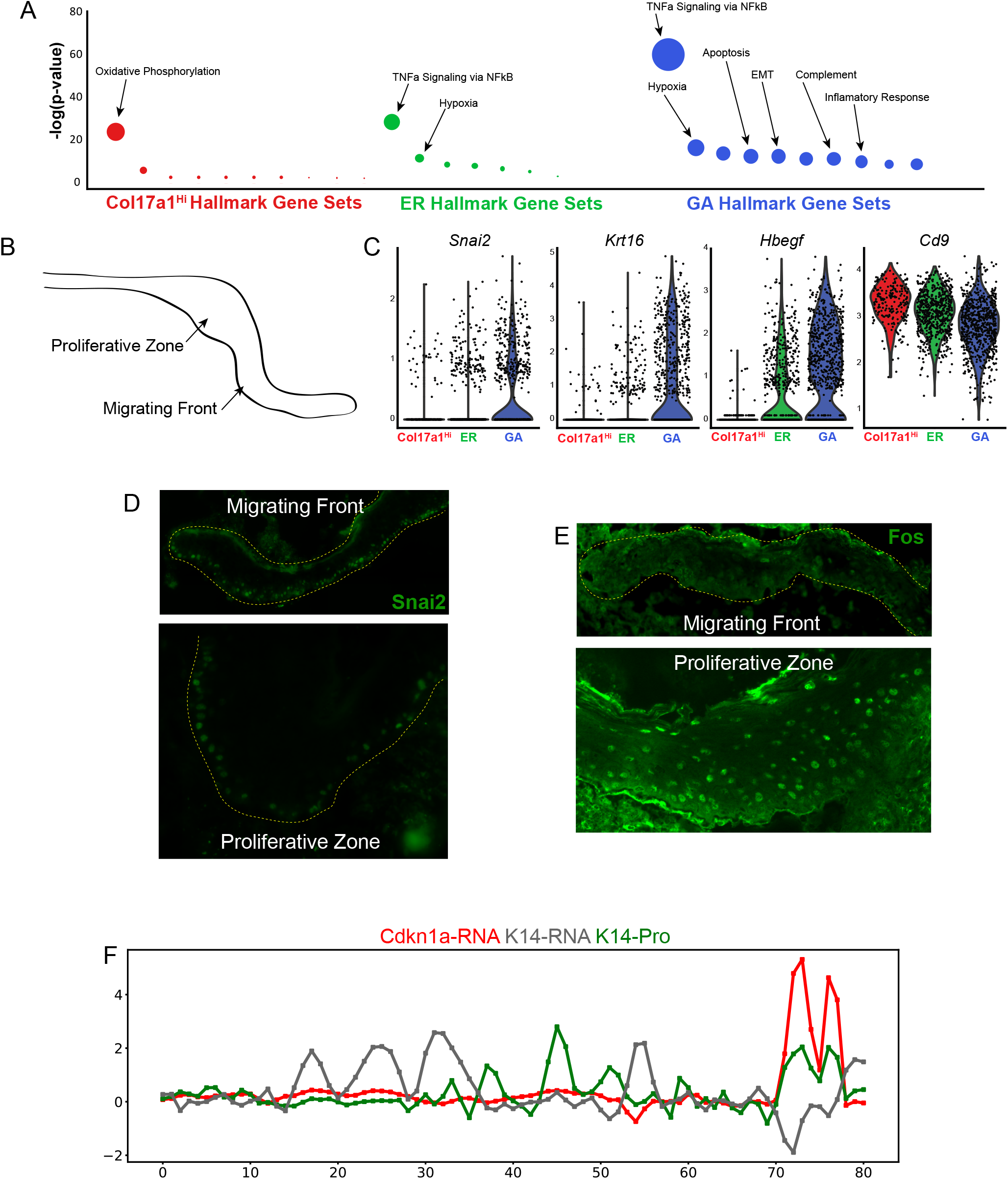
GO and marker gene analysis of basal cells form WO sample. A. GO analysis of WO basal cell subclusters. B. Diagram of equivalent wound with migrating front and proliferative zone. C. Known marker genes that have specific localization in epithelial cells around wound. D. Indirect immunofluorescence for Snai2 in different regions of wound. In the migrating front, the yellow dashed line outlines the entire migrating front. In the proliferative zone, the yellow dashed line marks the localization of the basement membrane. E. Indirect immunofluorescence for Fos in different regions of the wound. In the migrating front, the yellow dashed line outlines the entire migrating front. F. Local spatial autocorrelation analysis of RNAScope analysis of *Cdkn1a*, *Krt14*, and K14.

**Supplementary Figure 10:**
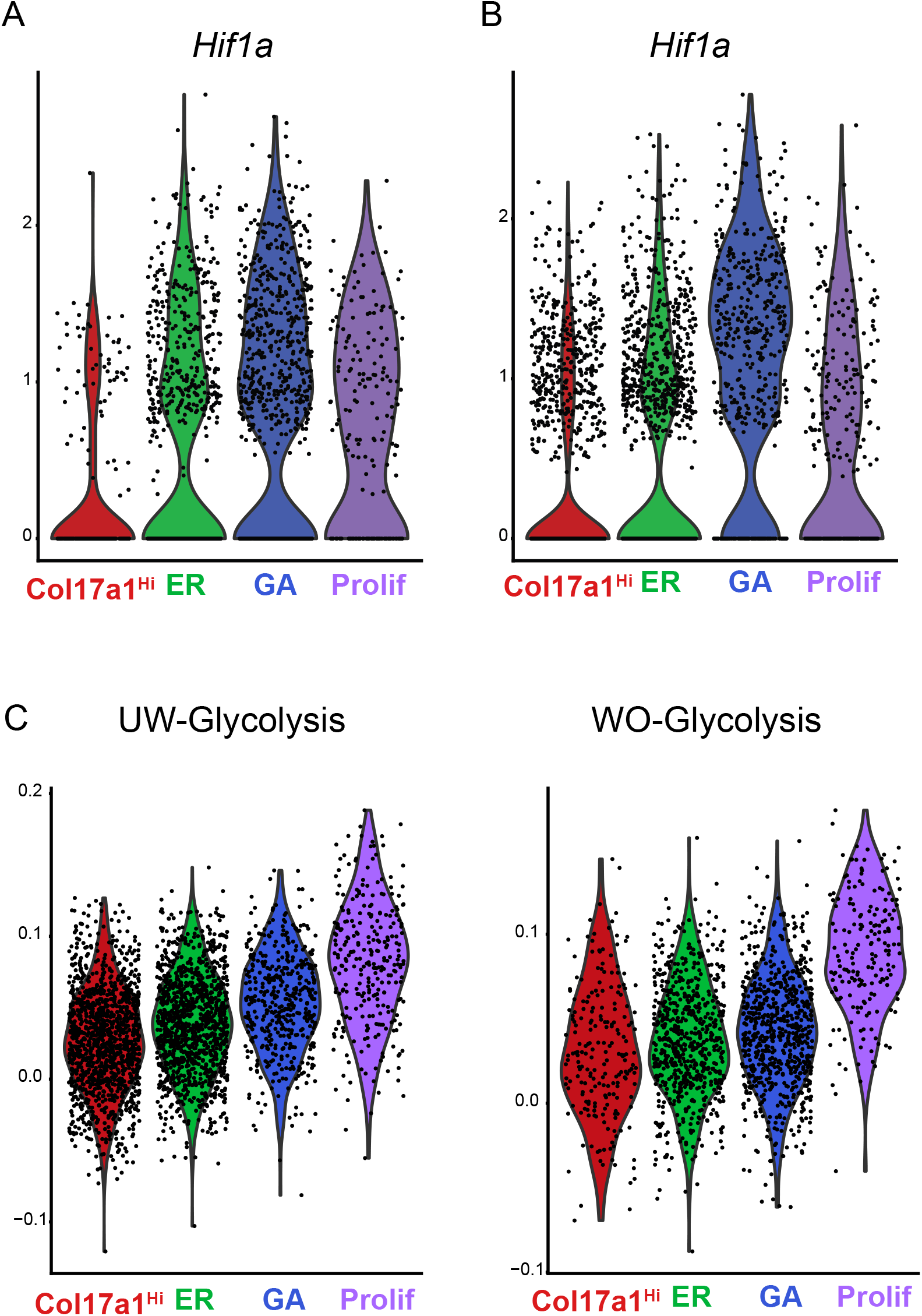
Scoring UW and UW basal cells for expression of metabolic genes. A. Violin plot for *Hif1a* expression in basal cells from WO sample. B. Violin plot for *Hif1a* expression in basal cells from the UW sample. C. Gene scoring analysis of all 4 basal subclusters from UW and WO samples using a glycolysis signature.

**Supplementary Figure 11:**
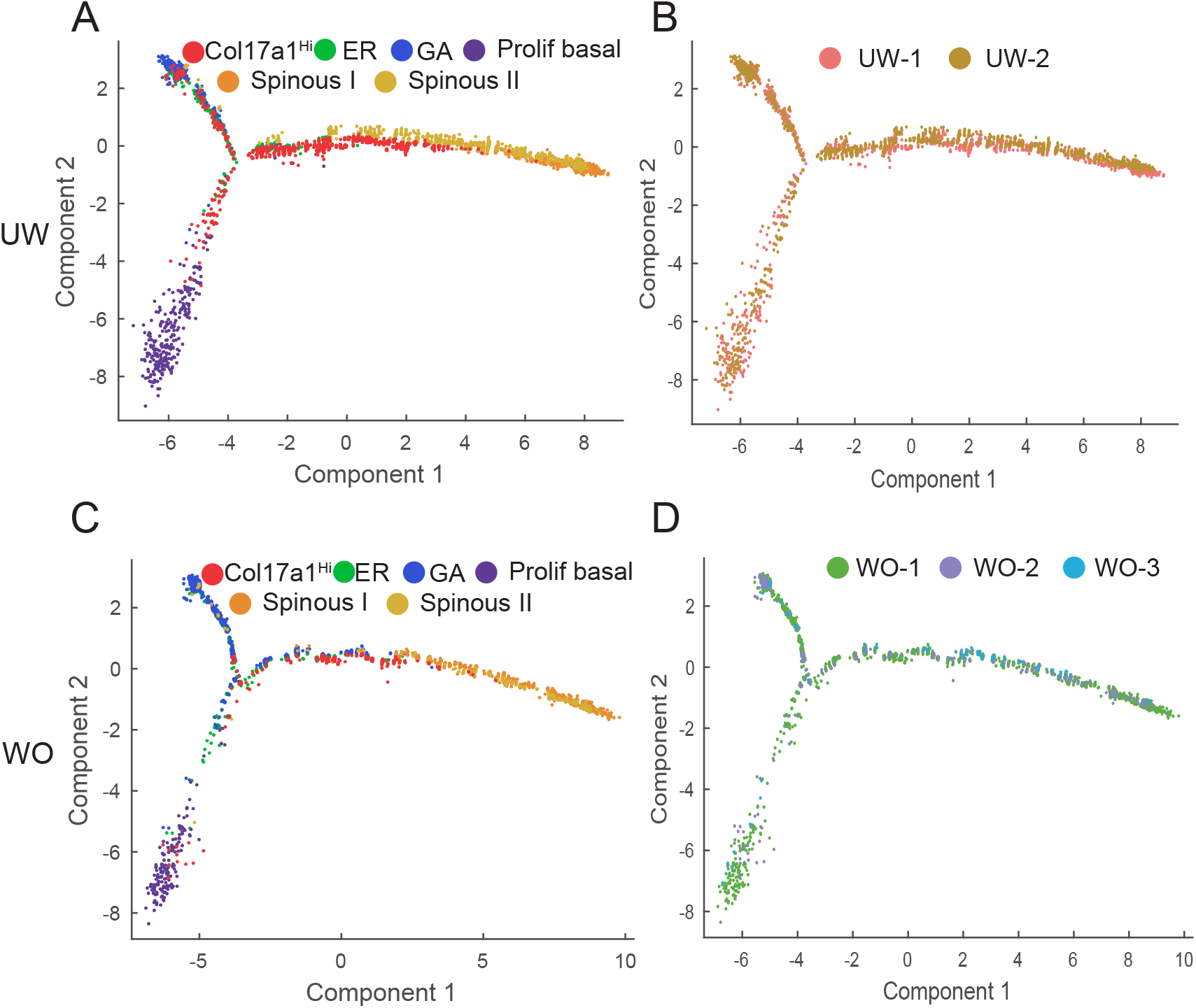
Lineage analysis by Monocle 2 in UW and WO skin. A. Lineage analysis of basal cells (proliferative and non-proliferative) and spinous cells in UW sample using Monocle 2. B. Monocle 2 analysis as in (A) but colored by sample replicate. C. Lineage analysis of basal cells (proliferative and non-proliferative) and spinous cells in WO sample using Monocle 2. D. Monocle 2 analysis as in (D) but colored by sample replicate.

**Supplementary Figure 12:**
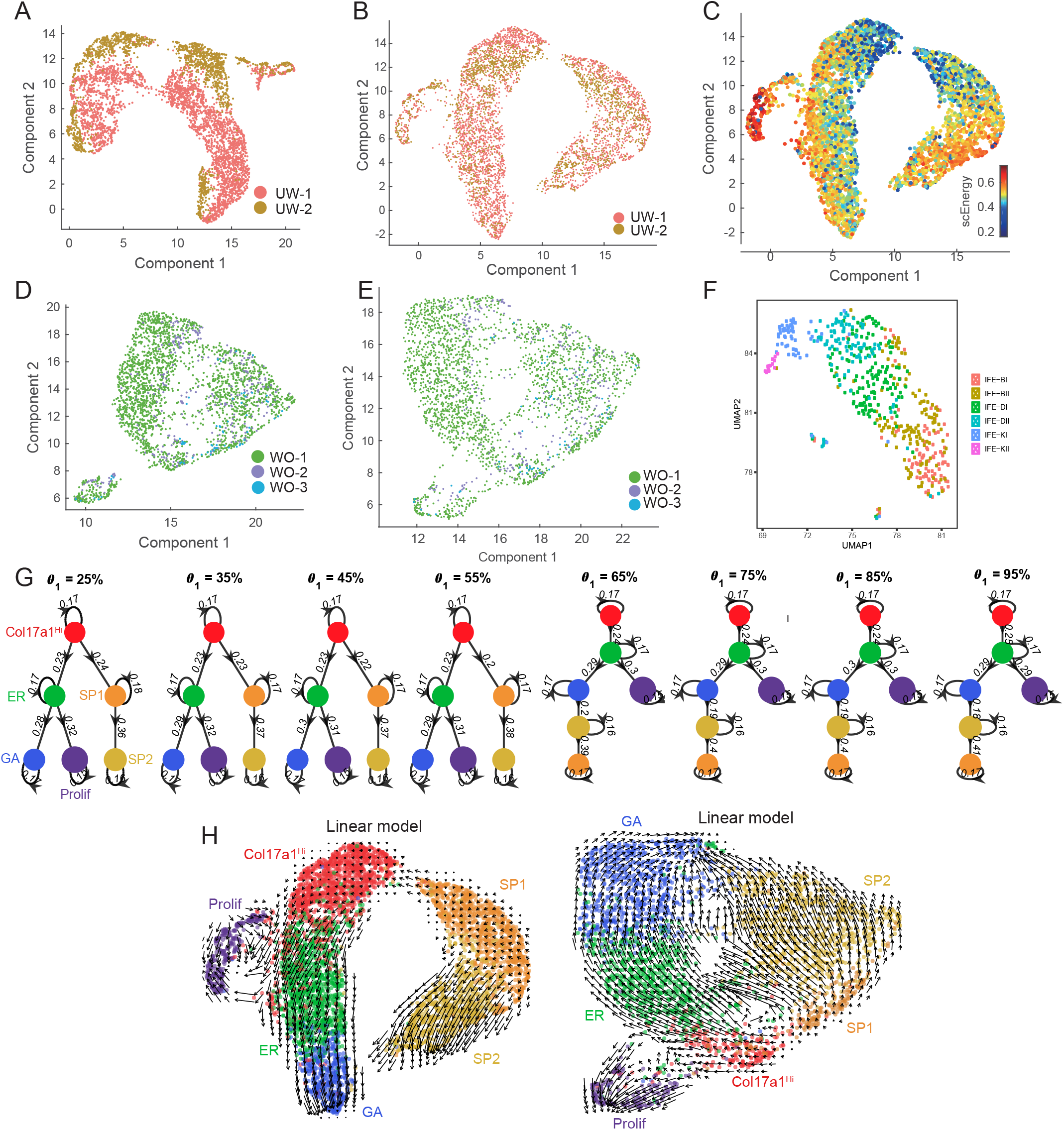
Lineage analysis by scEpath and RNA Velocity. A. Initial UMAP dimensional reduction for UW sample with cells identified by their replicate identity without batch correction. B. UMAP dimensional reduction for UW sample with cells identified by their replicate identity after batch correction. C. UMAP dimensional reduction for UW sample similar to (B) with scEnergy indicated. D. Initial UMAP dimensional reduction for WO sample with cells identified by their replicate identity without batch correction. E. UMAP dimensional reduction for WO sample with cells identified by their replicate identity after batch correction. F. UMAP dimensional reduction for 536 interfolicular epidermal cells from *Joost et al., 2016.* Cell color corresponds to exact annotations found in original paper. G. Inferred cell lineages of UW samples with different choices of parameter θ1 in scEpath. H. RNA velocity analysis using a linear model with vectors projected onto the batch corrected UMAP space for UW. I. RNA velocity analysis using a linear model with vectors projected onto the batch corrected UMAP space for WO.

**Supplementary Figure 13:**
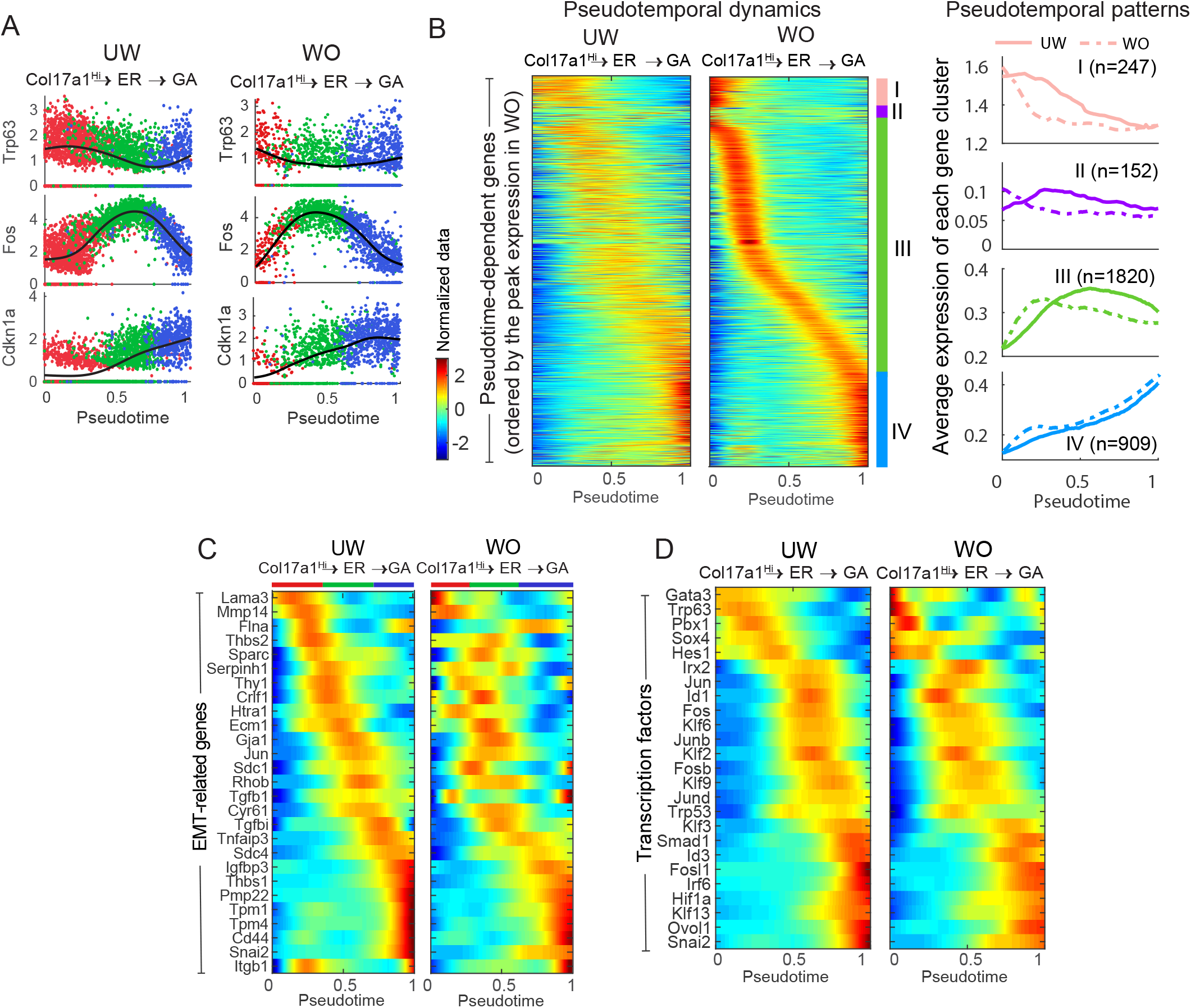
Pseudotemporal changes in gene expression. A. Dynamic changes in gene expression of individual genes *Trp63*, *Fos*, and *Cdkn1a* over pseudotime from the *Col17a1*^Hi^ to GA basal paths. B. Pseudotemporal dynamics of the identified 3,128 pseudotime-dependent genes (from WO sample) along the *Col17a1*^Hi^ to GA path in the UW and WO samples. Cell state identity (*Col17a1*^Hi^, ER or GA) was indicated on the top of each heatmap generated by the smoothed, normalized gene expression (for the colormap, blue and red colors indicate the low and high expression, respectively). Each row/gene was normalized to its peak value along the pseudotime. Distinct gene subclusters during pseudotime represented by pink, purple, green, and blue bars. The average gene expression of each gene subclusters was shown for UW (solid line) and WO (dashed line) samples, respectively. The number of genes in each gene subcluster was indicated. C. Subsets of EMT-related genes (from the 3,699 UW pseudotime-dependent genes) with their corresponding psudotemporal dynamics in both UW and WO basal cells along the *Col17a1*^Hi^ to GA path. D. Subsets of TFs (from the 3,699 UW pseudotime-dependent genes) with their corresponding psudotemporal dynamics in both UW and WO basal cells along the *Col17a1*^Hi^ to GA path.

## SUPPLEMENTARY TABLES

**Table S1:** Marker genes of each cluster from the UW sample.

**Table S2:** Marker genes of each cluster from the WO sample.

**Table S3:** Marker genes of each cluster from the epithelial cells of the UW sample.

**Table S4:** Marker genes of each cluster from the epithelial cells of the WO sample.

**Table S5:** Differentially expressed genes between basal cells from the UW and WO samples.

**Table S6:** Marker genes of the three basal subclusters from the UW sample.

**Table S7:** Marker genes of the three basal subclusters from the WO sample.

**Table S8:** Gene lists used for gene scoring.

